# COmapper: High-resolution mapping of meiotic crossovers by long-read sequencing in *Arabidopsis*

**DOI:** 10.1101/2024.10.21.619347

**Authors:** Dohwan Byun, Namil Son, Heejin Kim, Jaeil Kim, Jihye Park, Sang-jun Park, Hyein Kim, Jaebeom Kim, Juhyun Kim, Seula Lee, Youbong Hyun, Piotr A. Ziolkowski, Ian R. Henderson, Kyuha Choi

**Affiliations:** Department of Life Sciences, Pohang University of Science and Technology, Pohang, Gyeongbuk, Republic of Korea; School of Biological Sciences, Seoul National University, Seoul, Republic of Korea; Plant Quarantine Technology Center, Department of Plant Quarantine, Animal and Plant Quarantine Agency, Gimcheon, Republic of Korea; Laboratory of Genome Biology, Institute of Molecular Biology and Biotechnology, Adam Mickiewicz University, Poznań, Poland; Department of Plant Sciences, University of Cambridge, Cambridge, UK

**Author notes:** These authors contributed equally to this work (Dohwan Byun, Namil Son).

**Keywords:** Crossover, Long-read sequencing, Meiosis, RECQ4, Polymorphism

## Abstract

- Meiotic crossovers rearrange existing genetic variation between homologous chromosomes, profoundly affecting genomic diversity. Crossovers are typically constrained to 1–3 events per chromosome pair, and their distribution is shaped by chromatin accessibility and DNA polymorphisms. Genome-wide crossover maps can be generated in plants by high-throughput short-read sequencing or linked-read sequencing.
- Here, we use long-read nanopore sequencing technology to develop a crossover mapping pipeline, COmapper, for high-resolution mapping of genome-wide crossovers from pooled DNA of F1 hybrid pollen and F2 recombinant seedlings derived from a cross between Arabidopsis thaliana accessions Col and Ler. We validate the high accuracy of COmapper by applying nanopore long-read sequencing to pooled DNA of Arabidopsis F2 individuals with crossovers mapped by short-read sequencing.
- Using the COmapper, we constructed high-resolution genomic maps of crossovers using F_1_ hybrid pollen and F_2_ seedlings in wild type and crossover-elevated *recq4a recq4b* mutant, showing results comparable to short-read sequencing. Crossovers were enriched at gene-proximal promoters in wild type and increased but reshaped by high polymorphism density in *recq4a recq4b*.
- We propose that COmapper will be widely applicable for exploring the effects of genetic, epigenetic and environmental changes on the crossover patterns across diverse plant species.

## Introduction

Meiosis consists of a single round of DNA replication and two consecutive rounds of cell division, halving genome ploidy and producing genetically diverse gametes (Villeneuve & Hillers, 2001). During meiosis, homologous chromosomes undergo programmed DNA double-strand breaks (DSBs), pairing, synapsis and homology-directed repair, resulting in crossovers or non-crossovers (Hunter, 2015; Zickler & Kleckner, 2023). Topoisomerase-like SPO11 (SPORULATION) proteins catalyze the formation of meiotic DSBs and remain covalently attached to the ends of these DSBs (Grelon *et al*., 2001; Keeney *et al*., 2014; Lam & Keeney, 2014; Robert *et al*., 2016). Upon removal of the bound SPO11-oligonucleotides, the 5′ ends of DSBs undergo bidirectional 5′ to 3′ resection, producing 3′ overhangs of single-stranded DNA (ssDNA) (Neale *et al*., 2005; Lam & Keeney, 2014). This ssDNA invades non-sister chromatids and searches for homology, leading to the formation of recombination intermediates such as displacement loops and double Holliday junctions. Recombination intermediates are eventually resolved into either crossovers or non-crossovers (Mercier *et al*., 2015; Gray & Cohen, 2016; Wang & Copenhaver, 2018; Zickler & Kleckner, 2023). Although substantial DSBs are initially formed, only a few are repaired into crossovers by class I and class II repair pathways in most species. In plants, the class I pathway is responsible for ∼85–90% of all crossovers, which are formed by a group of pro-crossover factors including the E3 ligase HEI10 (HUMAN ENHANCER OF CELL INVASION NO.10) (Mercier *et al*., 2015; Wang & Copenhaver, 2018). Class I crossovers are sensitive to interference, a phenomenon that results in multiple crossovers being more widely spaced than would occur randomly. Crossover interference in plants is explained by a coarsening model where the diffusible and aggregating dynamics of HEI10 proteins limit closely spaced crossovers, resulting in 1–3 crossovers per pair of homologous chromosomes (Mercier *et al*., 2015; Morgan *et al*., 2021; Girard *et al*., 2023). In the class II pathway, ∼10–15% of all crossovers are non-interfering and mediated by the endonuclease MUS81 (MMS AND UV SENSITIVE 81) (Berchowitz *et al*., 2007; Zakharyevich *et al*., 2012). Class II crossovers are inhibited by different pathways involving anti-recombination factors, such as RECQ4A and RECQ4B helicases in plants (Crismani *et al*., 2012; Séguéla-Arnaud *et al*., 2015; Mieulet *et al*., 2018).

In plants, the distribution and frequency of meiotic DSBs and crossovers along chromosomes are non-random and depend on chromatin accessibility (Wijnker *et al*., 2013; Choi *et al*., 2013, 2018; Shilo *et al*., 2015; He *et al*., 2017; Underwood *et al*., 2018). Consequently, meiotic crossovers predominantly occur in gene-rich euchromatic regions and are suppressed in heterochromatic pericentromeres and centromeres. At a finer scale, crossovers are highly enriched at gene transcription start sites (TSS) near nucleosome-depleted promoters, but are rare in heterochromatic transposable elements and repeats (Wijnker *et al*., 2013; Choi *et al*., 2013; Choi & Henderson, 2015). In line with this, DNA methylation and histone H3 methylation on lysine 9 (H3K9me2) inhibit crossover formation (Yelina *et al*., 2015; Underwood *et al*., 2017, 2018; Underwood & Choi, 2019; Fernandes *et al*., 2024). Moreover, a high density of DNA polymorphisms and structural variation suppress crossovers (Choi *et al*., 2016; Serra *et al*., 2018a; Dluzewska *et al*., 2023; Fernandes *et al*., 2024). However, the frequency of meiotic crossovers correlates with the heterozygosity along chromosomes at a kilobase-(kb) or megabase-pair (Mb) scale in *Arabidopsis* (*Arabidopsis thaliana*), which depends on MSH2, a component of the mismatch repair system (Ziolkowski *et al*., 2015; Dluzewska *et al*., 2023). Therefore, mapping of genome-wide crossovers at high resolution is important for understanding the genetic and epigenetic effects on meiotic crossovers and genome evolution in various species (Henderson & Bomblies, 2021; Naish *et al*., 2021; Kim & Choi, 2022). Short-read sequencing and a probabilistic method for genotyping and crossover prediction have been intensively used to determine crossover numbers and distribution in F_2_ individuals between *Arabidopsis* accessions (Wijnker *et al*., 2013; Rowan *et al*., 2015; Serra *et al*., 2018b; Lawrence *et al*., 2019; Nageswaran *et al*., 2021; Kim *et al*., 2022, 2024). Recently, linked-read sequencing has been applied to DNA pooled from hybrid pollen grains or individual pollen nuclei of hybrid plants using microfluidics on a 10x Chromium Controller to map crossovers in *Arabidopsis*, tomato (*Solanum lycopersicum*) and beak sedge (*Rhynchospora breviuscula*) (Sun *et al*., 2019; Rommel Fuentes *et al*., 2020; Castellani *et al*., 2024).

Here, we develop a crossover mapping pipeline, COmapper (https://github.com/KyuhaChoi-Lab/COmapper), using long-read nanopore sequencing of pooled DNA from F_1_ pollen or F_2_ recombinant seedlings from *Arabidopsis* hybrids derived from a cross between the accessions Columbia-0 (Col) and Landsberg *erecta*-0 (L*er*). We validate the precision of COmapper for genome-wide crossover analysis through long-read nanopore sequencing of F_2_ recombinant individuals whose crossover sites were mapped by short-read sequencing. We demonstrate that COmapper allows high-resolution mapping of crossovers using long-read sequencing of pooled genomic DNA from wild-type and *recq4a recq4b* (*recq4ab*) F_1_ hybrid pollen grains or from their F_2_ recombinant seedlings. We propose that COmapper will be useful for determining crossover patterns at high resolution in diverse plants with different genetic backgrounds and ploidy.

## Materials and Methods

### Plant materials

The *Arabidopsis* accessions Col and L*er* were used as the wild types for Col × L*er* hybrid control plants. F_1_ and F_2_ hybrid plants were cultivated in a growth room under the controlled conditions of 20°C, 50–60% relative humidity and 16-h light/8-h dark photoperiod. The *recq4ab* mutants in the Col and L*er* backgrounds were used as previously described (Fernandes *et al*., 2018). *J3pro:J3^G155R^* plant was used as described (Kim *et al*., 2024). The *Arabidopsis* accession Di-G (Dijon, NASC stock number: N28205) was obtained from the Arabidopsis Biological Resource Center.

### Genotyping by sequencing (GBS)

GBS libraries were prepared as previously described (Rowan *et al*., 2015, 2019; Nageswaran *et al*., 2021; Kim *et al*., 2024). Briefly, genomic DNA (gDNA) was extracted from two or three mature leaves per individual plant using the CTAB (1% [w/v] CTAB, 50 mM Tris-HCl pH 8.0, 0.7 M NaCl, 10 mM EDTA) method. Leaves were collected into 2-ml tubes containing four 3-mm stainless steel beads, frozen in liquid nitrogen and ground using a mixer mill (Retsch, MM400). Then, 500 μl of CTAB buffer was added to the ground tissue, mixed vigorously and incubated at 65°C for 30 min. Following a chloroform extraction and gDNA precipitation with the addition of isopropanol, the gDNA pellet was washed with 1 ml 70% (v/v) ethanol twice, air-dried and resuspended in nuclease-free water. 150 ng of gDNA was used to generate one barcoded sequencing library per plant. The barcoded libraries were then pooled and subjected to paired-end 150-bp sequencing using an Illumina HiSeq-X platform (Macrogen, Korea). The TIGER pipeline was used to analyze the sequencing data and identify crossover sites (Rowan *et al*., 2015).

### Preparation of genomic DNAs for nanopore long-read sequencing

To purify pollen grains from F_1_ hybrid plants and extract their genomic DNA, F_1_ plants were planted in soil and grown at 20°C under a 16-h light/8-h dark photoperiod. Whole inflorescences of these hybrid plants were collected and transferred to ice-cold 10% (w/v) sucrose in a 50-ml tubes. Flowers were homogenized with a blender (Tefal, BL3051KR), filtered through 80-μm nylon mesh, and centrifuged at 350*g* for 10 min at 4 °C. The supernatant was discarded and the pellet was resuspended in 10% (w/v) sucrose, filtered through a 40-μm cell strainer, and centrifuged at 100*g* for 10 min at 4°C. The purified pellet was frozen in liquid nitrogen and ground with a mortar and pestle. The pellet powder was resuspended in CTAB buffer. Proteinase K (NEB, P8107S) and RNase A (20 mg/ml, Roche, 10109142001) were added to a final concentration of 80 U/ml and 20 μg/ml, respectively. To check pollen disruption, a 1-μl sample was diluted in a 1:10 ratio with 10% (w/v) sucrose and observed under a stereomicroscope (Leica, M165 FC). Then, 500 μl of the resuspended pellet was transferred to a 1.5-ml tube, to which an equal volume of 25:24:1 (v/v/v) phenol:chloroform:isoamylalcohol (Sigma, 77617) was added, and the mixture was homogenized by gentle manual shaking. The tube was centrifuged at 15,000*g* for 10 min at 4°C, and 400 μl of supernatant was transferred into a new tube, to which a Nanobind disk (Nanobind® Plant Nuclei Kit RT, PacBio, 102-302-000) was added. The tube containing the Nanobind disk was gently mixed, 400 μl isopropanol was added, and the tube was slowly inverted and incubated for 2 h at room temperature on a rotator set at 7 rpm. A magnetic rack (DynaMag^TM^-2, Invitrogen, 12321D) was used to separate the disk and the supernatant, and the supernatant was removed. PW1 buffer in the Nanobind Plant Nuclei Kit was used for washing the disk twice and the supernatant was removed completely by centrifugation. Then, 150 μl EB+ buffer was added onto the disc and gently tabbed and incubated overnight at room temperature. The supernatant was transferred to a new 1.5-ml tube and stored at −20°C. The DNA was quantified using a Qubit dsDNA Broad Range assay kit (Thermo Fisher, Q32853) and integrity checked by gel electrophoresis.

For nanopore sequencing of gDNA from 1,000 pooled seedlings, freshly harvested F_2_ seeds from F_1_ hybrid plants were surface-sterilized, germinated, and grown on 1/2 MS agar medium supplemented with 1% sucrose under controlled conditions (20°C, 16-hour light/8-hour dark photoperiod). Seedlings were harvested 10 days after germination and immediately frozen in liquid nitrogen. The 10-day-old seedlings were powdered in liquid nitrogen using a pre-chilled mortar and pestle. gDNA was extracted using the CTAB method and quantified using a Qubit dsDNA Broad Range assay kit (Thermo Fisher, Q32853). Nine micrograms of gDNA from pollen or seedlings was used to construct a nanopore long-read sequencing library using a Ligation Sequencing Kit V14 (Nanopore, SQK-LSK114). The libraries were sequenced on a PromethION platform (BGI, Hong Kong).

### Alignment of nanopore sequencing reads to the reference genome

The FASTQ files containing the nanopore sequencing reads were aligned to the TAIR10 reference genome sequence using minimap2 with the following settings “-ax map-ont -cs“(Li, 2018). To obtain uniquely aligned long reads, sambamba was employed with the following settings: “-F “not unmapped and not duplicate and mapping_quality >= 30 and sequence_length >= 100““(Tarasov *et al*., 2015). Bases with a quality (QUAL) score <14 were replaced with ‘N’ using an awk script, thereby addressing genotyping errors that were a consequence of sequencing errors.

### COmapper pipeline

To obtain high-quality SNPs distinguishing the Col and L*er* accessions, we followed the previous method with modifications (Fernandes *et al*., 2024). SNPs within syntenic regions were identified by comparing the reference genomes of Col and L*er* using SyRI (Goel *et al*., 2019). These SNPs were validated using sequencing data from a total of 2,160 Col × L*er* F_2_ plants were used (Rowan *et al*., 2019). To retain only high-quality SNPs, the following criteria were applied: (1) genotyping results available for at least 40 F_2_ plants, (2) allele frequency between 0.4 and 0.6, and (3) adherence to Mendelian segregation across the 2,160 F_2_ individuals (chi-square test, *P*-value > 0.1). These sites were then integrated with an independent set of high-quality SNP calls from the previous report (Yelina *et al*., 2015).

The genotypes for the SNPs in masked alignments were converted to string format using a reference list of 226,864 high-quality Col/L*er* SNPs. Genotype strings with two successive identical genotypes at either end (CC-CC or LL-LL) were identified as parental reads. Genotype strings that satisfied the following conditions were identified as crossover reads: The two successive SNP genotypes at one end differed from those at the other end (CC-LL or LL-CC), and the four successive SNP genotypes before and after the crossover site matched the genotype at the respective end of the genotype string (CCCC-LLLL or LLLL-CCCC). Genotype strings with less than eight characters or those not identified as either crossover or parental reads were excluded from the calculation of crossover frequency. Regions exhibiting a depth of coverage exceeding the average genome read depth by three-fold and exhibiting a high frequency of false positives in the negative control provided by the sequencing of Col × L*er* F_1_ leaf gDNA were excluded from the analysis. The standard output provided the input alignment information, the genotyping summary, the total number of crossover and parental reads, the length of the crossover and parental reads and the information regarding crossover reads in a CSV file.

### Bioinformatic analysis of crossovers and genomic features

To analyze the genome-wide pattern of crossovers as well as genes, transposable elements, DNA methylation, SPO11-1–oligos, and MNase-seq (micrococcal nuclease digestion with deep sequencing) data, the TAIR10 genome was tiled into 100-kb bins. DNA methylation levels in each bin were used for plotting. The coverage of SPO11-1–oligo and MNase-seq data was normalized by sequencing reads of randomly digested gDNA before plotting (Choi *et al*., 2018). The number of overlaps for genes, transposable elements, and crossovers within each bin was calculated and plotted. The coverage of SPO11-1–oligo and MNase-seq data and the frequency of crossovers around genes were calculated using deepTools computeMatrix (v. 3.5.4) to generate gene metaplots. Regions within the pericentromeric regions and where the distance between consecutive SNPs was >4,000 bp were excluded from all analyses to increase resolution of crossover sites. For the analysis of SNP density around crossover site, the SNP frequency in 10-bp bins within a 4-kb window centered on each crossover site was calculated using deepTools computeMatrix (v. 3.5.4).

### Accession numbers

Arabidopsis Genome Initiative locus identifiers for the genes mentioned in this article are as follows: AT3G13170 for SPO11-1, AT1G10930 for RECQ4A, AT1G60930 for RECQ4B, AT1G53490 for HEI10, AT4G30870 for MUS81, AT3G18524 for MSH2, and AT3G44110 for J3.

## Results

### Establishment of COmapper and long-read sequencing to map crossovers

To develop COmapper and enable high-resolution mapping of meiotic crossovers at the single-nucleotide polymorphism (SNP) level using long-read sequencing, we used the two polymorphic *Arabidopsis* accessions, Columbia-0 (Col) and Landsberg *erecta*-0 (L*er*) (Fig. **1a**). We crossed them to generate wild-type (WT) Col × L*er* F_1_ hybrid plants, from which we collected recombinant pollen; we also allowed them to self-pollinate to collect F_2_ recombinant seeds (Fig. **1a**). We performed genotyping by sequencing (GBS) of individual F_2_ recombinant plants using individual indexing, short-read sequencing and the TIGER pipeline, which allows the mapping of crossover sites along chromosomes (Fig. **1b**) (Wijnker *et al*., 2013; Rowan *et al*., 2015, 2019; Nageswaran *et al*., 2021; Kim *et al*., 2022, 2024). We then used the GBS data and pooled genomic DNA (gDNA) of F_2_ recombinant seedlings as a positive control to establish and validate COmapper. Notably, GBS of F_2_ recombinants has been also used to validate the linked-sequencing-based crossover mapping method via microfluidics on the 10x Chromium Controller (Fig. **1c**) (Sun *et al*., 2019). GBS and linked sequencing both allow the genotyping of individual genomes or large DNA molecules through individual barcoding and multiplexing, amplification, short-read sequencing, demultiplexing and read alignment to determine crossover sites (Fig. **1b,c**). By contrast, nanopore long-read sequencing followed by COmapper can directly identify Col-specific or L*er*-specific SNPs within individually sequenced DNA molecules without prior indexing or amplification and can determine the position of crossover sites in individual DNA molecules (Fig. **1d,e**). Therefore, the read length obtained by long-read sequencing and the number of SNPs in individually sequenced DNA molecules are critical for generating genome-wide high-resolution maps of crossovers, along with the depth of genome sequence coverage.

**Fig. 1.**
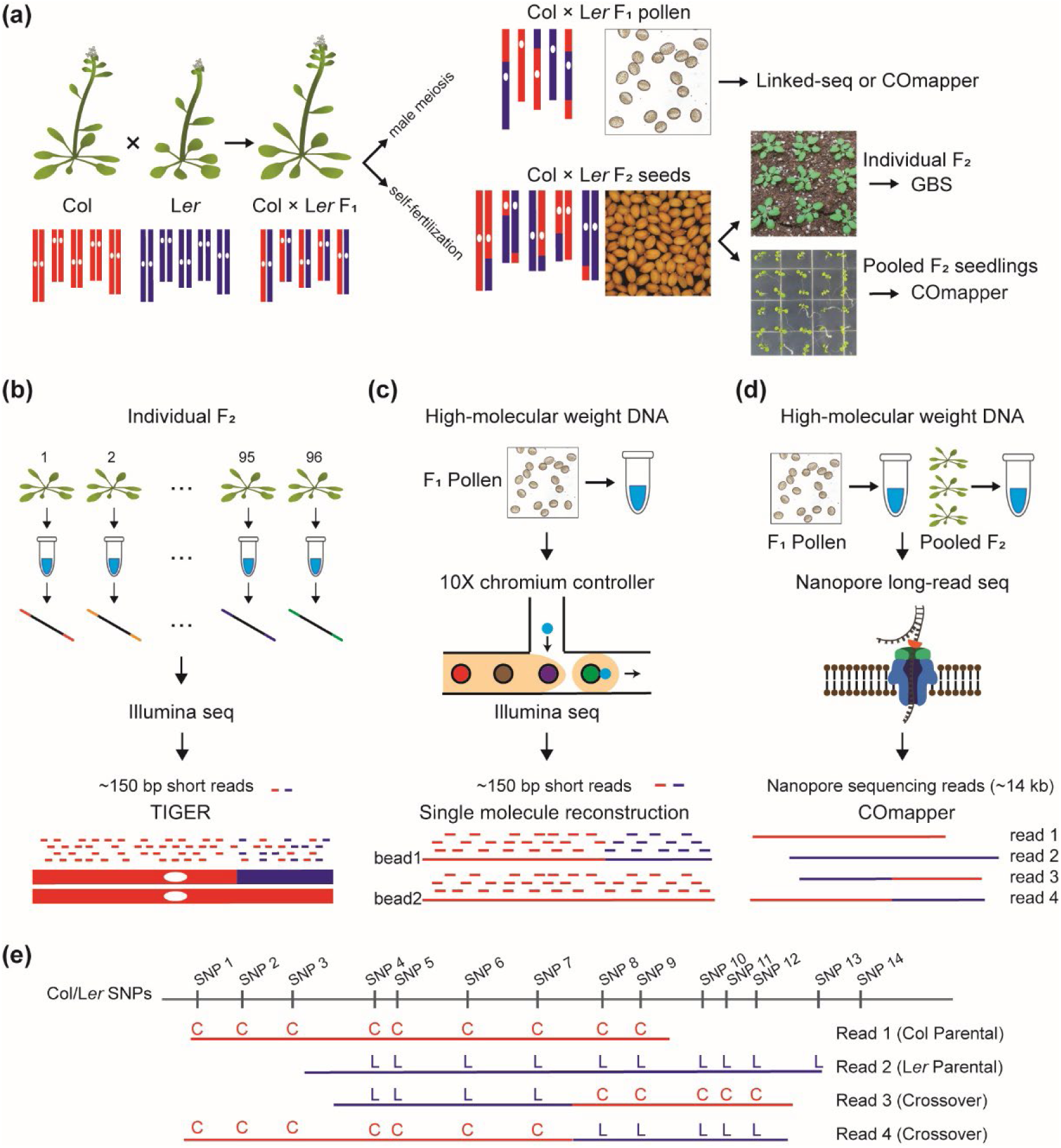
Genome-wide mapping of crossovers using long-read nanopore sequencing. (**a**) iagrams illustrating the preparation of recombinant pollen from F_1_ hybrids and F_2_ plants to map crossovers. (**b**) Mapping of genome-wide crossovers in F_2_ plants by genotyping-by-sequencing (GBS) using multiplex barcoding and the TIGER pipeline. (**c**) Mapping of crossovers in pollen DNA from F_1_ hybrid plants using 10x Chromium Controller and linked-read sequencing. (**d**) Mapping of crossovers in F_1_ pollen and F_2_ plants using nanopore long-read sequencing and COmapper. (**e**) High-resolution mapping of crossover sites by analyzing single-nucleotide polymorphisms (SNPs) within long-read DNA molecules. Red C indicates a Columbia-0 (Col)-specific SNP; purple L indicates Landsberg *erecta*-0 (L*er*) SNP.

To obtain long-read nanopore sequencing reads for the mapping of crossover sites (Fig. **2a**), we isolated mature pollen grains from WT Col × L*er* F_1_ hybrid plants and extracted large genomic DNA (gDNA) molecules using our modified CTAB method (see Methods). Our gDNA extraction from pollen grains of Col × L*er* F_1_ hybrids resulted in nanopore sequence read lengths of ∼11.1 kb for gDNA (mean, 11.1 kb; N50, 14.0 kb), comparable to that of gDNA extracted from F_2_ seedlings using the CTAB method (Figs **2b****, S1**). To achieve deep sequencing coverage, we used ∼9 µg gDNA on the PromethION device, which produced reads corresponding to ∼100–150 Gb of sequence per flow cell (Fig. **S1**; Table **S1**). We successfully aligned ∼90.7% of the nanopore reads to the TAIR10 *Arabidopsis* reference genome using the minimap2 alignment tool (Fig. **2c**). The aligned reads covered the genome uniformly from each chromosome end to end except across centromere-proximal regions, with a mean read coverage frequency of 904.9 (Figs **2d****, S2**). We used a high-quality SNP dataset distinguishing between Col and L*er* accessions previously used for GBS (Rowan *et al*., 2015; Serra *et al*., 2018b; Kim *et al*., 2024), with a 524-bp mean distance between consecutive SNPs (Fig. **2e,f**). We noted that structural variation between Col and L*er* results in decreased coverage of the aligned long reads and fewer SNPs for GBS in the pericentromeric regions near centromeres (Figs **2d,e****, S3**). Of all aligned long reads, 86.1% contained more than eight SNPs per read, with a mean 19.8 SNPs per read (Fig. **2g**), which were used to determine crossover sites by COmapper.

**Fig. 2.**
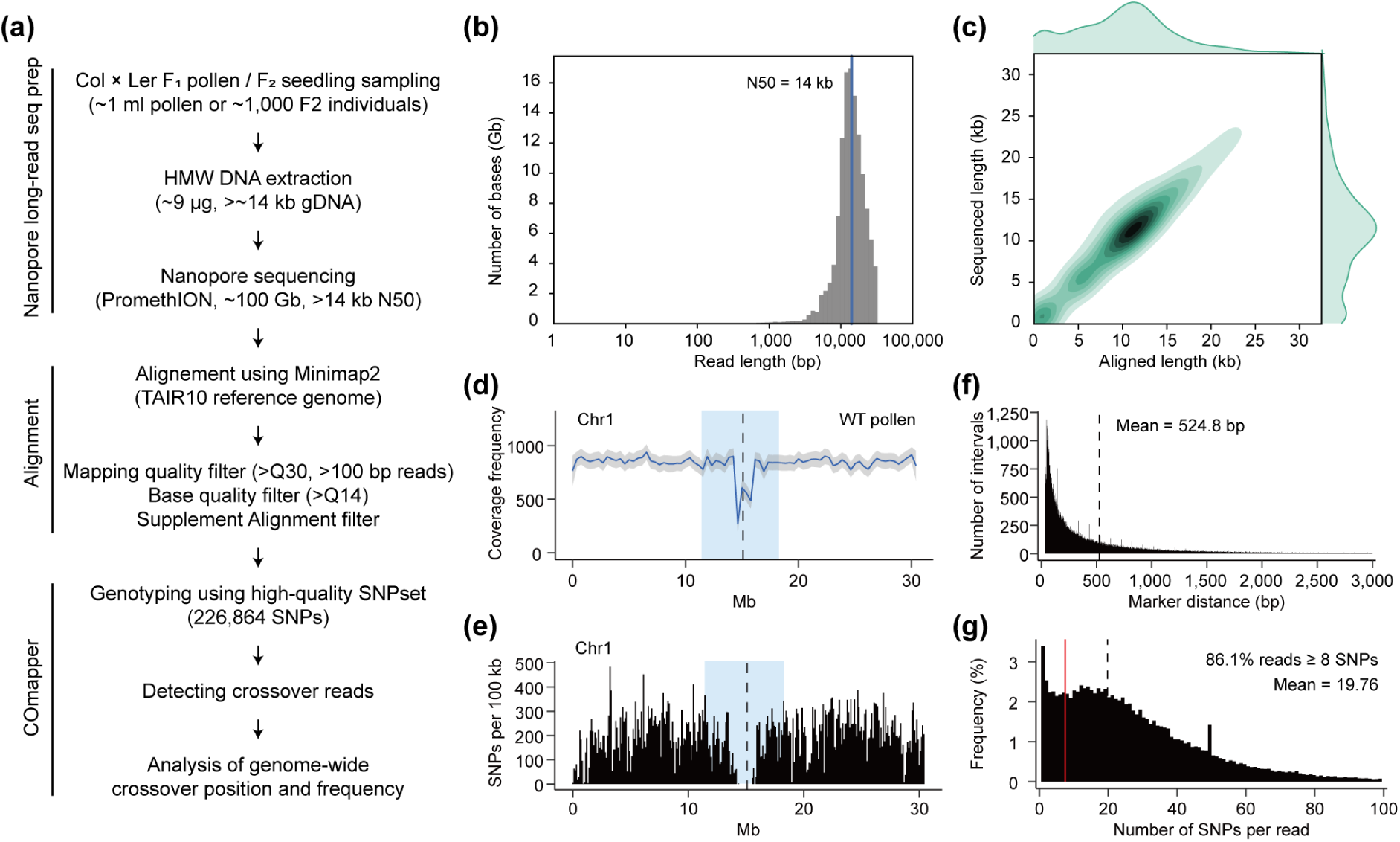
Development of COmapper to map crossovers using long-read sequencing. (**a**) Procedure for crossover mapping using nanopore long-read sequencing and the COmapper pipeline. (**b**) Distribution of read length and number of pollen DNAs sequenced by nanopore long-read sequencing. (**c**) Density distribution of the lengths of DNA molecules sequenced by nanopore long-read sequencing and aligned to the *Arabidopsis* reference genome using Minimap2. (**d**) Coverage frequency of long reads from Col × L*er* F_1_ pollen DNA along *Arabidopsis* chromosome 1. The vertical dashed line indicates the centromere. The pericentromeric region is highlighted in blue. (**e**) As in (**d**), but showing the density of SNPs between the Col and L*er* accessions, in 100-kb sliding windows. (**f**) Distribution of inter-SNP distances between consecutive SNPs differentiating the Col and L*er* accessions. The vertical dashed line indicates the mean distance between consecutive SNPs. (**g**) Distribution of SNP number and frequency of SNPs per nanopore sequencing read. The vertical dashed line indicates the mean number of SNPs per read, and the vertical red line indicates the position corresponding to eight SNPs in a read.

### Development and validation of COmapper

We sought to establish a crossover mapping model for COmapper, which scans for the presence of SNPs from the 5′ to the 3′ end of a long read and identifies simple crossovers (SCOs) where consecutive Col-specific SNPs switch to consecutive L*er*-specific SNPs, or vice versa, by counting the numbers of SNPs (Fig. **3a**). In *Arabidopsis*, approximately 50% of crossovers are associated with gene conversion (Wijnker *et al*., 2013), which are detected in COmapper data as SCOs characterized by a single switch between parental SNPs. However, a subset of crossovers contain two or more switches between parental SNPs, representing complex crossover (CCOs) (Schweiger *et al*., 2024). Therefore, we also considered CCOs when identifying crossover sites by examining the number and positions of consecutive Col-specific or L*er*-specific SNPs (Fig. **3b**). Accordingly, we developed and tested six crossover mapping models (Fig. **3b**), which use different numbers of consecutive Col-specific and L*er*-specific SNPs with or without multiple switches between parental SNPs within the crossover tract to map crossovers. To validate these models, we performed nanopore sequencing of two independent pooled gDNA samples from 4 or 46 Col × L*er* F_2_ recombinant plants in which crossover sites had already been determined by GBS (Fig. **S4**; Table **S2**). We then examined the precision of each crossover mapping model by aligning the long-read molecules to the *Arabidopsis* genome and analyzing their SNPs. This revealed that the 8-SCO and 8-CO models were the most accurate in detecting the true crossover sites mapped by GBS, with precisions of 97.3% (8-SCO) and 97.4% (8-CO), respectively (Welch′s *t*-test; 8-SCO vs 8-CO *P* = 0.747, 8-CO vs 7-CO *P* = 1.38× 10^−11^) (Figs **3c–e**). In contrast, 4-SCO model showed a high number of false positives, likely due to the inherent per-base sequencing error rate (3.98%, Q score = 14) of nanopore reads (Fig. **S5**). The 8-CO model, which identifies both SCO and CCO types, detected more crossovers than the 8-SCO model, which detected only the SCO type (Welch′s *t*-test, *P* = 2.82 × 10^−10^) (Figs **3c,d****, S6,7**). COmapper with the 8-CO model detected ∼7.3% and ∼1.5% of the CCO type of total crossover sites from the pooled gDNA of 4 and 46 F_2_ plants, respectively (Figs **3f****, S6,7**). Furthermore, using COmapper with the 8-CO model, we identified crossover sites in nanopore reads of pooled hybrid pollen and pooled F_2_ seedlings of wild-type (WT) Col × L*er* and *recq4ab* Col × *recq4ab* L*er*. Among detected crossovers, ∼8.9% (WT) and ∼7.8% (*recq4ab*) were CCOs in F_1_ hybrid pollen. In F_2_ seedlings (*n* = 1,000), CCOs accounted for 2.1% (WT) and 2.9% (*recq4ab*) similar to the 2.1% observed to 46 F_2_ plants analyzed by GBS and COmapper (Fig. **3f**).

**Fig. 3.**
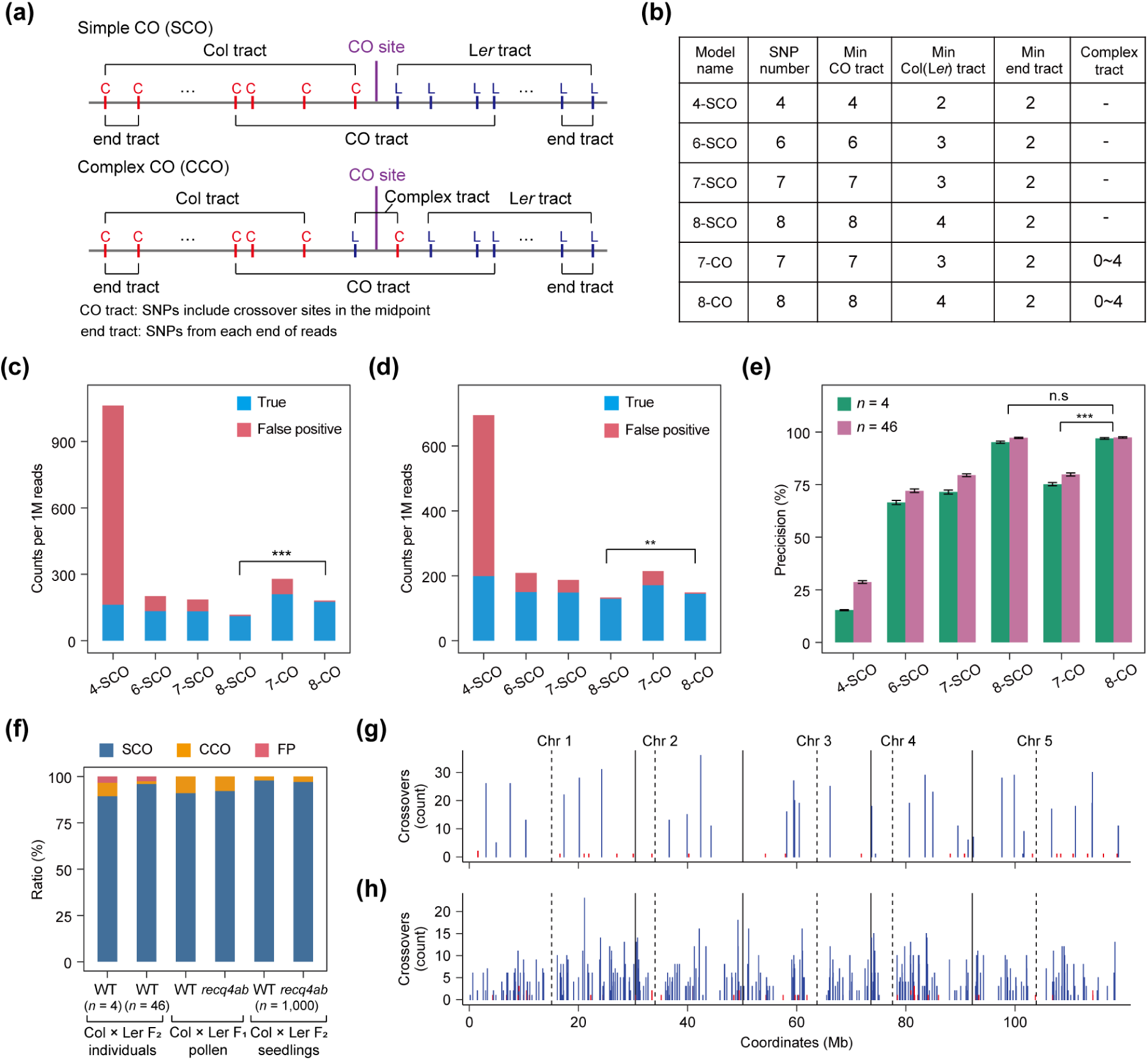
Validation of COmapper using Col × L*er* F_2_ recombinant plants. (**a**) Diagram showing simple crossover (SCO) and complex (CCO) types. (**b**) Summary table of the tested COmapper models with different numbers of SNPs and minimum crossover (Min CO) and complex tracts. (**c**) Number of crossovers detected per million reads in different COmapper models in 4 Col × L*er* F_2_ plants (Kim *et al*., 2024). True crossovers (blue) were detected by COmapper models among the crossovers mapped by GBS in 4 F_2_ plants, and false crossovers (red) indicate the crossovers that differ from the crossovers defined by GBS. (**d**) as in (**c**), but showing number of crossovers detected per million reads in 46 Col × L*er* F_2_ plants. (**e**) Precision (True positive/(True positive + False positive)) of crossover detection in the different COmapper models shown in (C). In (C) and (D), asterisks indicate significant differences between models (\**P* < 0.05, \*\**P* < 0.01, \*\*\**P* < 0.001; Welch′s *t*-test). (**f**) Relative proportion of crossover types in 4 F_2_ plants (genotyped by GBS), 46 F_2_ plants (genotyped by GBS), Col-0 × L*er* F_1_ hybrid pollen DNA, *recq4ab* Col × *recq4ab* L*er* hybrid pollen DNA, 1,000 Col-0 × L*er* F_2_ seedlings and 1,000 *recq4ab* Col × *recq4ab* L*er* F_2_ seedlings detected by the established model (8-CO) of COmapper. FP (red) indicates the proportion of false positive crossovers identified by COmapper in 4 and 46 F_2_ plants compared to the crossovers identified by GBS. (**g**) Frequency of crossovers mapped by COmapper along the five *Arabidopsis* chromosomes on a continuous *x*-axis in 4 Col × L*er* F_2_ recombinants, where crossovers in each F_2_ plant were mapped by GBS. Blue bars represent true positive crossovers mapped by GBS; red bars represent false positive crossovers. (**h**) As in (**g**), but from 46 Col × L*er* F_2_ recombinants.

To validate the output of COmapper, we used the crossover sites mapped by GBS and TIGER with manual curation as true positives from 4 and 46 Col × L*er* F_2_ recombinant plants (33 crossovers for 4 F_2_ plants, 342 crossovers for 46 F_2_ plants) (Dataset **S1** and **S2**). COmapper with the 8-CO model was capable of identifying true positive crossover sites multiple times per site, whereas ∼3% false positives were detected only once or twice (Fig. **3g,h**). Analysis of these false positives revealed four sources: chimeric reads with error clusters (16 in 4 F_2_, 16 in 46 F_2_), substitution errors (5 in 4 F_2_, 12 in 46 F_2_), chimeric reads with large indels (3 in 46 F_2_), and mitotic recombination (10 in 46 F_2_), reflecting limitations of nanopore sequencing (Fig. **S8**; Dataset **S3**).

COmapper accurately identified all 33 crossovers that were mapped by GBS in the 4 F_2_ recombinants, with a mean of 18.9 long reads covering each crossover site when reaching a sequencing depth of 40.14 Gb (Fig. **S4a**,**b**). Of the 342 crossovers mapped by GBS in the 46 F_2_ recombinants, COmapper accurately mapped 299 crossovers, with a mean of 6.07 long reads per crossover site. Notably, COmapper failed to detect the remaining 43 crossovers. To analyze the false negatives, we categorized the detected (299) and undetected crossovers (43) from the 46 F_2_ pool according to the 8-SNP length around the crossover site (Fig. **S9a,b**). While crossovers were more concentrated in the SNP-rich regions, the proportion of undetected crossovers relative to the total crossover (detected and undetected crossovers) increases as the 8-SNP length increases (Fig. **S9a,b**). It indicates the dependance of COmapper’s crossover mapping sensitivity on SNP density. However, the undetected crossovers (false negatives) were not concentrated at certain region of genome but randomly distributed at genome-wide scale (Fig. **S9c**; Dataset **S4**).

As COmapper depends on the depth of sequencing, we conducted a comprehensive analysis of how sequencing coverage influences COmapper’s performance in terms of true positive (TP) and false positive (FP) crossover read detection, precision (TP / [TP + FP]), and false negatives (crossovers detected by GBS but not by COmapper) (Fig. **S10**). We found that both TP and FP reads, including those from the same crossover sites, increase proportionally with sequencing depth, keeping precision stable across coverage levels (Fig. **S10a–f**). As coverage per F_2_ increases, false negatives decrease. At the same time, the discovery rate of new unique crossovers declines due to repeated detection of previously identified sites (Fig. **S10g,h**). Additionally, we found that pooling more individuals increases the likelihood of capturing non-overlapping crossover sites, thereby reducing redundancy and maximizing the number of distinct crossovers (Fig. **S10i**). This validation shows that SNP-resolution crossover mapping by COmapper requires long reads with at least eight SNPs and sufficient sequencing depth (Table **S1**). Taken together, these results confirm that our COmapper pipeline with an 8-CO model (hereafter referred to simply as ‘COmapper’) can directly map crossovers in long-read nanopore sequencing molecules at SNP resolution with an overall precision of 97%.

To further assess COmapper’s performance in the *recq4ab* background, we re-sequenced genomic DNA from 39 *recq4ab* Col × L*er* F_2_ individuals—previously analyzed by GBS—using nanopore sequencing at a total coverage of 410× (∼10.5×per individual) (Fig. **S11**). COmapper achieved ∼96% precision, comparable to the 97% observed in the WT dataset, demonstrating reliable crossover detection in both genotypes. In addition, we confirmed that COmapper is highly robust to Col-CEN (Telomere-to-Telomere) reference genome as well as TAIR10 in crossover detection using 46 F_2_ individuals (Fig. **S12**).

### Mapping of crossovers from a pool of Col × L*er* hybrid pollen grains by COmapper

Next, we used the COmapper and long-read nanopore sequencing technology to directly map crossovers from pooled gDNA samples of WT Col × L*er* and *recq4ab* Col × *recq4ab* L*er* F_1_ hybrid pollen grains (Fig. **4**). Our GBS analysis showed a 3.12-fold increase in crossovers during male meiosis in the *recq4ab* mutant compared to the WT, consistent with the previous report (Fig. **4a**) (Fernandes *et al*., 2018). Similarly, COmapper detected a 2.41-fold increase in crossovers per genome (119 Mb) when analyzing long-read sequencing data produced from *recq4ab* F_1_ hybrid pollen DNA (mean = 3.92 crossovers per genome, ∼763.8x coverage) compared to the WT (mean = 1.62 crossovers per genome, 1,072.3x coverage) (Welch′s *t*-test, *P* = 1.22 × 10^−12^) (Fig. **4b**). Importantly, COmapper rarely detected crossovers as false positive crossover events in the negative control sample (leaf genomic DNA from Col × L*er* F_1_ hybrids; mean = 0.03 crossovers per genome, 82.8x coverage) compared to the WT Col × L*er* F_1_ hybrid pollen (Welch′s *t*-test, *P* = 5.51 × 10^−10^) (Fig. **4b**; Table **S1**). These results indicate that COmapper enables the detection of meiotic crossovers in pooled gDNA from Col × L*er* hybrid pollen grains, with an increased number of crossovers in a *recq4ab* background.

**Fig. 4.**
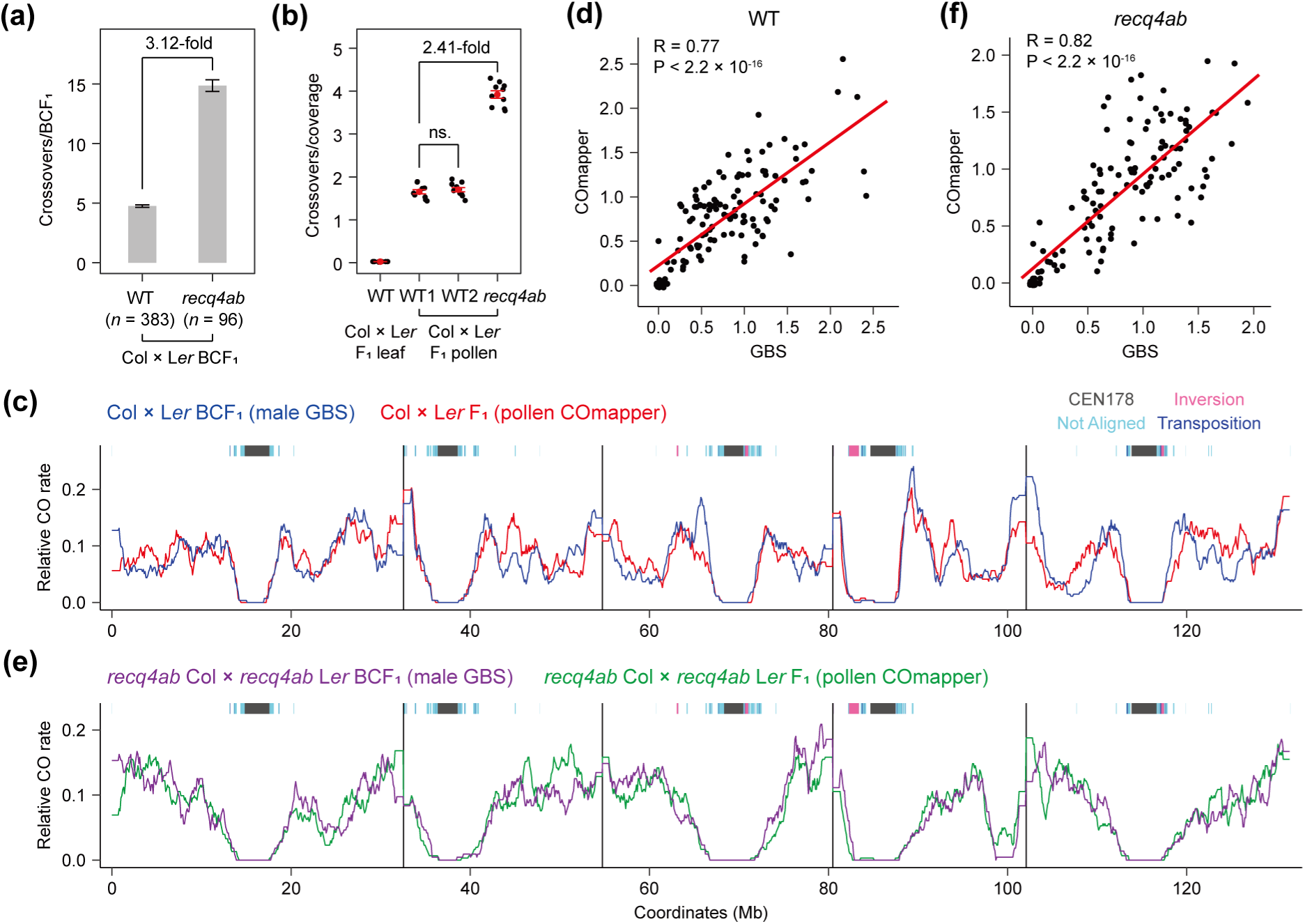
Mapping of crossovers from pooled F_1_ hybrid pollen DNA by COmapper. (**a**) Number of male crossovers per BCF_1_ individual calculated from GBS data of WT Col × L*er* BCF_1_ (*n* = 383) (Kim *et al*., 2024) and *recq4ab* Col × *recq4ab* L*er* BCF_1_ (*n* = 96) plants. *n* indicates the number of backcrossed F_1_ plants. Data are presented as means ± SD. (**b**) Number of crossovers per genomic coverage of nanopore sequencing data of gDNA from WT Col × L*er* F_1_ leaves, WT Col × L*er* F_1_ pollen (WT set1, WT set2) and *recq4ab* Col × *recq4ab* L*er* F_1_ pollen. Each data point indicates crossovers per genome from a randomly selected million reads. Data are presented as means ± SD from 10 replicates. (**c**) Relative frequency of crossovers mapped by Col × L*er* male GBS (blue, *n* = 1,824 crossovers) and Col × L*er* F_1_ pollen COmapper (red, *n* = 1,679 crossovers) along the five *Arabidopsis* chromosomes on a continuous *x*-axis in Col × L*er* male meiosis using Col-CEN complete genome. Fisher′s exact test was applied to each 100-kb bin in every 1 Mb window to assess differences in relative crossover rates. (**d**) Correlation between crossover frequencies in 1-Mb bins of Col × L*er* BCF_1_ individuals (measured by GBS) and Col × L*er* F_1_ pollen (measured by COmapper) across the genome, assessed using Pearson′s correlation test. (**e**) As in (**c**), but showing the relative crossover frequency of crossovers mapped by *recq4ab* Col × *recq4ab* L*er* male GBS (blue, *n* = 1,428 crossovers) and *recq4ab* Col × *recq4ab* L*er* F_1_ pollen COmapper (red, 1,963 crossovers). (**f**) As in (**d**), but showing the correlation between crossover frequencies of crossovers mapped by *recq4ab* Col × *recq4ab* L*er* BCF_1_ individuals and *recq4ab* Col × *recq4ab* L*er* F_1_ pollen.

Previously, GBS-based crossover maps of *Arabidopsis* Col × L*er* male meiosis using a backcross (BC) population detected ∼4.47 crossovers per genome (Kim *et al*., 2024). In comparison, COmapper detected ∼1.65 crossovers per genome in nanopore long-read sequencing of pooled pollen grains from Col × L*er* hybrid plants, with all detected 1,204 crossovers being non-overlapping. This suggests an inherent feature of COmapper method, which samples crossover molecules directly from long reads, rather than reconstructing full meiotic products, as is done in GBS. To further validate COmapper’s performance, we performed an independent COmapper experiment using pollen from WT Col × L*er* F_1_ plants, confirming the robustness of its crossover detection (Figs **4b****, S13**). Despite detecting fewer absolute events, our validation demonstrates that COmapper provide a consistent and representative subset of crossovers relative to sequencing coverage.

To compare genome-wide crossover patterns detected by COmapper and GBS using the Col-CEN complete genome (Kim *et al*., 2024), we calculated relative crossover rates as the proportion of crossovers in each 100-kb genomic bin, normalized to the total number of crossovers (Fig. **4c–f**). We used Fisher’s exact test to assess crossover frequency differences and found no significant regional differences between COmapper and GBS, in either WT or *recq4ab* datasets (Fig. **4c,e**). We also observed that the genomic landscape of crossovers mapped by COmapper correlated with that of GBS (Fig. **4d,f**), indicating that COmapper and GBS produce highly comparable genome-wide crossover landscapes. Notably, we identified a ∼3.2 Mb inversion (Chr4: 14,564,462–17,818,793) in *recq4ab* via nanopore long-read analysis, which likely underlies crossover suppression observed in both GBS and COmapper, consistent with the previous report (Serra *et al*., 2018b). Inversions interfere with homolog pairing and synapsis, and single crossovers within them can produce dicentric or acentric chromosomes, leading to suppression or rare double crossovers (Rönspies *et al*., 2022). Although not statistically significant, COmapper detected slightly more crossovers within the *recq4ab* inversion (Fig. **4e**) than GBS, possibly due to selection against non-viable recombinants in the F_2_ population.

Despite the comparable crossover landscapes generated by both approaches, we noted that COmapper detects fewer crossovers at sub-telomeric regions than GBS by plotting the crossovers of male meiosis along a telomere-to-centromere axis that combined all five chromosomes (Fig. **S14a**). The mean SNP density was lower at sub-telomeric regions than in mid-chromosomal regions (Fig. **S14b,c**). Additionally, the physical distances covered by the eight SNPs required for crossover-site mapping by COmapper were shorter than those identified by GBS in the sub-telomeric, mid-chromosomal and pericentromeric regions (Fig. **S14d–f**).

To further assess the effect of SNP density on the crossover detection sensitivity of COmapper, we divided the genome into four categories based on the SNP density, defined by the 8-SNP span (Fig. **S15a,b**), and compared the distribution of COmapper-detected crossovers across these regions with those of GBS-detected crossovers (Fig. **S15c**). In the WT male GBS dataset, ∼14.6% of crossovers were found in pericentromeric and subtelomeric regions with 8-SNP spans ≥10 kb, in contrast, only 1.8% of crossovers in WT pollen COmapper dataset were detected in these SNP-poor regions (Fig. **S15c**). Since GBS-based crossover detection is not affected by SNP density, this result indicates that COmapper has reduced sensitivity in SNP-poor regions and preferentially detects crossovers in SNP-rich areas. However, because crossovers in SNP-poor pericentromeric regions account for only a small fraction of total crossovers, and a subset of SNP-poor regions are interspersed among SNP-rich regions. (Fig. **S15a**), the overall effect of SNP density on crossover rate profiles along chromosomes was minimal when analyzed at the megabase scale. Nonetheless, comparisons of crossover rates between COmapper and GBS should be interpreted cautiously in SNP-poor regions, such as subtelomeric and pericentromeric regions, particularly at high resolution (e.g., kilobase scale), where detection sensitivity may differ signficantly. COmapper requires long reads (≥10–15 kb) containing at least eight SNPs for reliable crossover detection, underscoring its limitation in regions with spare SNP density. Despite this, we demonstrate that COmapper can effectively map genome-wide crossovers and capture the genetic effect of the *recq4ab* mutant on crossover frequency and distribution by nanopore sequencing of pooled hybrid pollen DNA.

### Mapping of crossovers from pooled F_2_ recombinant seedlings by COmapper

GBS analysis of Col × L*er* F_2_ individuals revealed ∼8 crossovers per F_2_ (Serra *et al*., 2018b; Fernandes *et al*., 2018; Rowan *et al*., 2019; Nageswaran *et al*., 2021; Kim *et al*., 2022, 2024). From the COmapper analysis of 40 Gb nanopore sequencing data from the 4 F_2_ pool and 100 Gb data from the 46 F_2_ pool, the proportions of non-overlapping crossovers relative to the total detected crossovers were 5.23% (33/630) and 18.02% (299/1659), respectively (Fig. **S10**). Thus, we sought to increase the proportion of detected non-overlapping crossovers by increasing the number of F_2_ individuals included in the pool. To determine pool size, we conducted subsampling analysis using 4 F_2_ pool and fitted generalized linear model (GLM) (*y*∼−0.196 *ln*(*x*) + 0.873) (Fig. **S10i**). Based on the COmapper results from the 46 WT F_2_ pool, approximately 1,600 crossovers are expected to be detected from 100 Gb of sequencing data. According to our GLM model, sequencing 1,000 F_2_ individuals at a total depth of 100 Gb—equivalent to ∼0.82× coverage per F_2_—would yield 90.2% non-overlapping crossovers, corresponding to 1,472 unique crossover events.

To test this hypothesis, we performed nanopore long-read sequencing of gDNA pooled from 1,000 Col × L*er* F_2_ recombinant seedlings (10 days old) derived from WT Col × L*er* and *recq4ab* Col × *recq4ab* L*er* F_1_ hybrid seedlings, and mapped their crossovers with COmapper (Fig. **5**). Much as the GBS analysis of Col × L*er* and *recq4ab* Col × *recq4ab* L*er* F_2_ individuals (30-day-old plants) detecting a 3.67-fold increase in crossover numbers in the *recq4ab* background compared to WT Col × L*er* F_2_ plants (Fig. **5a**) (Serra *et al*., 2018b; Kim *et al*., 2024), COmapper detected 3.86-fold more crossovers in the *recq4ab* Col × *recq4ab* L*er* F_2_ seedlings compared to WT Col × L*er* F_2_ seedlings (Fig. **5b**). COmapper generated crossover landscapes from the pooled F_2_ seedlings (*n* = 1,000) similar to those obtained by GBS from adult F_2_ individuals in both WT and *recq4ab* backgrounds, with crossovers highly enriched along chromosome arms and suppressed in pericentromeric and centromeric regions (Fig. **5c**–**f**). Notably, COmapper detected lower crossover frequencies in specific subtelomeric regions of chromosome 1 and 3 in *recq4ab* compared to WT (Fig. **5e**). Collectively, these results suggest that COmapper can be applied to a pool of hybrid F_2_ progeny as well as to F_1_ pollen for genome-wide mapping of crossovers and distribution in *Arabidopsis*.

**Fig. 5.**
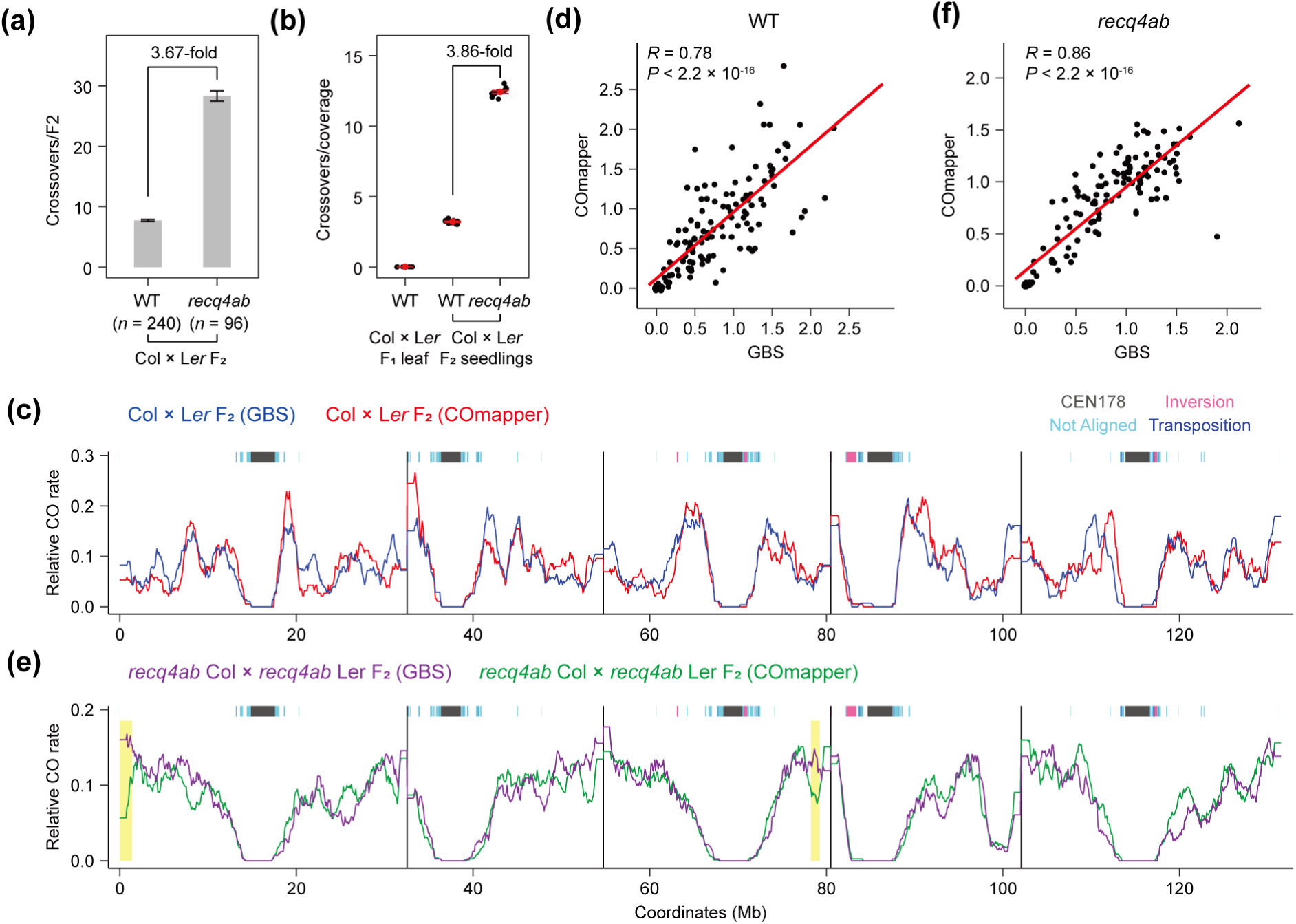
Mapping of crossovers from pooled F_2_ recombinant seedlings by COmapper. (**a**) Number of crossovers per F_2_ individual calculated from GBS data of WT Col × L*er* F_2_ (*n* = 240) (Kim *et al*., 2024) and *recq4ab* Col × *recq4ab* L*er* F_2_ (*n* = 96) plants. *n* indicates the number of F_2_ individual plants. Data are presented as means ± SD. (**b**) Number of crossovers per genome of nanopore sequencing data from gDNA of Col × L*er* F_1_ leaves, Col × L*er* F_2_ and *recq4ab* Col × *recq4ab* L*er* F_2_ seedlings (*n* = 1,000). Each data point indicates the number of crossovers per genome from a randomly selected 1 million reads. Data are presented as means ± SD from 10 replicates. (**c**) Relative frequency of crossovers mapped by GBS (blue, *n* = 1,852 crossovers) and COmapper (red, *n* = 1,390 crossovers) along the five *Arabidopsis* chromosomes on a continuous *x*-axis in WT Col × L*er* F_2_ plants (Col-CEN). Fisher′s exact test was applied to each 100-kb bin to assess differences in relative crossover rates. Bins with false discovery rate (FDR) below 0.05 are highlighted in yellow. (**d**) Correlation between crossover frequencies in 1-Mb bins of Col × L*er* F_2_ individuals (measured by GBS) and Col × L*er* F_2_ seedlings (measured by COmapper) across the genome, assessed using Pearson′s correlation test. (**e**) As in (**c**), but showing the relative frequency of crossovers mapped by GBS in *recq4ab* Col × *recq4ab* L*er* F_2_ plants (purple, *n* = 2,731 crossovers) or by COmapper in *recq4ab* Col × *recq4ab* L*er* F_2_ seedlings (green, *n* = 5,760 crossovers). (**f**) As in (**d**), but showing the correlation between crossover frequencies of crossovers mapped by *recq4ab* Col × *recq4ab* L*er* F_2_ individuals and *recq4ab* Col × *recq4ab* L*er* F_2_ seedlings.

### High-resolution mapping of meiotic crossovers around genes

In *Arabidopsis*, genome-wide meiotic DSB profiling by sequencing SPO11-1– oligonucleotides (SPO11-1–oligos) revealed that meiotic DSBs occur predominantly in nucleosome-depleted regions, with the highest peak observed over gene-proximal promoters, while the formation of DSBs at the exons of protein-coding genes with high nucleosome density is suppressed (Fig. **6a**) (Underwood *et al*., 2018; Choi *et al*., 2018). In agreement with this result, meiotic crossovers in plants are highly enriched around gene TSS (Hellsten *et al*., 2013; Wijnker *et al*., 2013; Choi *et al*., 2013; Choi & Henderson, 2015; Shilo *et al*., 2015; Kianian *et al*., 2018). Because we generated high-resolution crossover maps using COmapper, we examined the position and frequency of crossovers around the TSS and transcription end sites (TES) of all genes in a 4-kb window (2 kb on either side of the TSS or TES). Like meiotic DSBs that were previously determined by sequencing of SPO11-1–oligos (Choi *et al*., 2018), the crossovers mapped by COmapper in gDNA pooled from WT Col × L*er* F_1_ hybrid pollen showed higher enrichment in gene-proximal promoters but lower enrichment in gene bodies, compared to the same number of random sites (Figs **6a,b****, S16a,b**). The pattern of crossovers determined by our GBS dataset, which had ∼5.2x genome coverage for male meiosis (Kim *et al*., 2024), showed a similar enrichment around gene TSS as that mapped by COmapper in Col × L*er* F_1_ hybrid pollen pool (Figs **6b****, S16b**). Furthermore, COmapper and GBS with ∼4.5x genome coverage both detect a higher enrichment of crossovers around gene-proximal promoters and a lower enrichment in gene bodies in WT Col × L*er* F_2_ recombinant seedlings compared to random (Figs **6c****, S16c**), suggesting that chromatin accessibility is a major determinant in shaping crossovers as well as meiotic DSBs around genes in *Arabidopsis*.

**Fig. 6.**
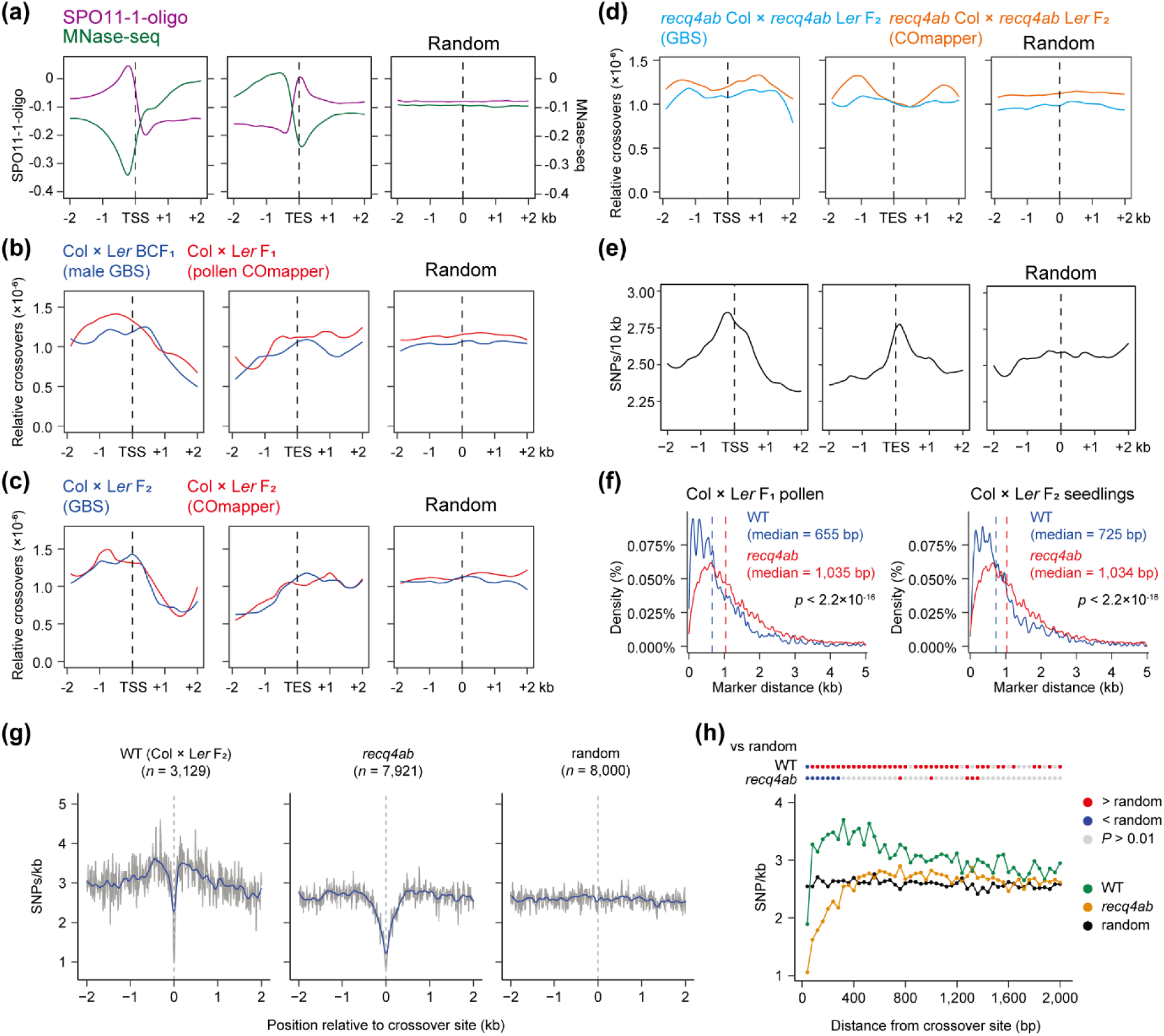
Analysis of crossovers and SNPs around genes. (**a**) Metaplots showing the density of SPO11-1–oligonucleotides (SPO11-1–oligo) and nucleosomes from MNase sequencing (MNase-seq) around gene transcription start sites (TSS) and transcription end sites (TES) in a 2-kb window (Choi *et al*., 2018). (**b**) As in (**a**), but showing the relative rate of crossovers mapped by GBS (blue, *n* = 1,824 crossovers) described (Rowan *et al*., 2019; Kim *et al*., 2024) and COmapper (red, *n* = 1,679 crossovers) in *Arabidopsis* Col × L*er* male meiosis. (**c**) As in (**b**), but showing the relative rate of crossovers mapped by GBS (blue, 1,852 crossovers) and COmapper (red, *n* = 1,391 crossovers) in *Arabidopsis* Col × L*er* F_2_ plants. (**d**) As in (**c**), but showing the relative rate of crossovers mapped by GBS (skyblue, *n* = 2,731 crossovers) and COmapper (orange, *n* = 5,760 crossovers) in *Arabidopsis recq4ab* Col × *recq4ab* L*er* F_2_ plants. (**e**) As for (**a**), but showing the density of SNPs between Col and L*er* accessions. (**f**) Density and physical distance between two consecutive SNPs at crossover sites in WT Col × L*er* and *recq4ab* Col × *recq4ab* L*er* for F_1_ pollen and F_2_ progeny. The vertical dashed line indicates median distance (bp) between two consecutive SNPs at the crossover site. Significance between genotypes was examined using a Wilcoxon test. (**g**) Metaplots showing SNP density around crossovers mapped by GBS and COmapper in WT Col × L*er* F_2_ and *recq4ab* Col × *recq4ab* L*er* F_2_ progeny compared to random positions. (**h**) As in (**g**), but showing the significance of changes in SNP density according to distance from the crossover site. Each dot represents the SNP density per 40-bp bin. Significance between genotype and a random position per interval was examined using a Wilcoxon test. Red and blue dots indicate higher and lower SNP density than random, respectively; gray dots indicate non-significant differences.

The GBS with sparse coverage (∼0.1–1x) and TIGER, a GBS pipeline for mapping crossovers are inexpensive and useful for measuring the number of crossovers and interference from many recombinant individuals (Rowan *et al*., 2015, 2019). However, when crossovers mapped by GBS and TIGER at different genome coverages (0–1×, 1–2×, and ≥2×) were plotted around TSS and TES at finer scales—based on the distance between the two SNPs flanking each crossover site—we found that high-resolution crossover mapping by GBS requires a genome coverage of at least 3×, which is comparable to COmapper enabling detection of crossovers at SNP resolution (Fig. **S17**).

### SNP density promotes interfering crossovers and inhibits non-interfering crossovers

We examined the effect of the *recq4ab* mutant on the pattern of crossovers mapped by COmapper or GBS. The loss of RECQ4A/B function resulted in a decreased crossover frequency at the gene TSS and TES, with an increase in crossovers at gene bodies (Figs **6d****, S16d**). Genetic disruption of *MSH2* was shown to further increase the frequency of class II crossovers in Col × L*er* F_2_ progeny in the *recq4ab* background, suggesting that a high density of DNA polymorphisms inhibits the formation of class II crossovers in *recq4ab* (Dluzewska *et al*., 2023). Consistent with this possibility, we detected peaks for high SNP density between the Col and L*er* accessions at gene TSS and TES, with lower SNP density at distal promoters, gene bodies and random sites (Fig. **6e**). We also found that the physical distance between consecutive SNPs at crossover sites is greater in *recq4ab* than in WT based on nanopore long-read sequencing of F_1_ pollen and F_2_ progeny (Fig. **6f**), suggesting that the increased class II crossovers in *recq4ab* are more sensitive to SNP-mediated crossover inhibition and tend to occur in regions with lower SNP density compared to class I crossovers in WT.

To further investigate the effect of SNP density on the distribution of class I and class II crossovers, we plotted the density of SNPs in a 2-kb window, upstream and downstream of the center positions of crossovers mapped by COmapper and GBS, in the WT Col × L*er* and *recq4ab* Col × *recq4ab* L*er* backgrounds (Fig. **6g**). The density of SNPs around the crossovers was higher in WT but lower in the *recq4ab* background compared to that at random sites, with the lowest density seen at the center of crossover positions in both WT and *recq4ab* (Fig. **6g,h**). We also found a similarly higher density of SNPs around crossover sites compared to random sites in the GBS analysis of F_2_ individuals from Col expressing the *J3^G155R^* allele (*J3pro:J3^G155R^*) × L*er* and Col × Di-G F_1_ hybrids (Fig. **S18**). *J3^G155R^*, a dominant negative allele of *J3* encoding a HSP40 co-chaperone, increases class I crossovers by enhancing the abundance of HEI10 (Kim *et al*., 2024). These results suggest that the class I crossovers occur preferentially in the regions with higher SNP density compared to random sites, whereas the increased class II crossovers due to the loss of RECQ4A/B function occur mainly in regions with lower SNP density. It is worth noting that the SNP density nearest to the crossover site is lower for both class I and class II crossovers, which is consistent with previous observations that high-density SNPs inhibit crossovers at the crossover hotspot scale (Borts & Haber, 1987; Serra *et al*., 2018a; Szymanska-Lejman *et al*., 2023). Taken together, these results indicate that COmapper allows the generation of high-resolution genome-wide crossover maps and reveals the effects of SNP density and genetic mutation of *RECQ4A/B* on the positioning and frequency of crossovers.

## Discussion

Next-generation and linked-read sequencing technologies have contributed to our understanding of the patterning of genome-wide meiotic crossovers in diverse species (Rowan *et al*., 2015; Serra *et al*., 2018b; Kianian *et al*., 2018; Sun *et al*., 2019; Nageswaran *et al*., 2021; Kim *et al*., 2022, 2024; Durand *et al*., 2022; Castellani *et al*., 2024). Here, using nanopore long-read sequencing (Wang *et al*., 2021), we developed COmapper that can generate high-resolution maps of meiotic crossovers from pooled pollen and F_2_ progeny of F_1_ hybrid plants (Figs. **3–6**). We demonstrated that COmapper can map crossovers along chromosomes at high resolution by validating the detection of crossovers in genomic DNA pooled from 4 or 46 recombinant F_2_ plants, in which crossovers per each plant were mapped using the GBS approach with individual barcoding.

At a chromosomal scale, COmapper generated genome-wide crossover landscapes comparable to those of GBS in WT and *recq4ab* F_1_ hybrid pollen and F_2_ seedlings from self-pollinated F_1_ hybrids, with high enrichment at chromosome arms and suppression at pericentromeres and centromeres (Figs. **4,5**). At a finer scale, COmapper revealed a strong enrichment of crossovers at gene promoters and a reduction within gene bodies, resembling the distribution of meiotic DSBs (Fig. **6a–c**). Analysis of SNP density around crossovers showed distinct patterns: it was positively correlated with class I crossovers but negatively with class II (Fig. **6g,h**), demonstrating differential sensitivity of the two crossover pathways to sequence polymorphisms (Ziolkowski *et al*., 2015; Blackwell *et al*., 2020; Dluzewska *et al*., 2023; Szymanska-Lejman *et al*., 2023).

We observed that COmapper can mis-map some crossovers occurring within SNP-poor genomic regions, because accurate crossover mapping depends on both SNP density and read length—each long read must cover a sufficient number of SNPs (Figs. **S14**). In genomes with low SNP density, alternative methods like GBS, linked-read sequencing or Hi-C may offer more reliable crossover detection. Like COmapper, Hi-C and linked-read approaches can be applied to pooled samples, but typically provide lower crossover resolution. GBS requires at least 3x coverage to achieve crossover detection resolution comparable to COmapper. In pooled samples, the number of crossovers detected by COmapper increased proportionally with nanopore sequencing depth (Table **S3**), indicating that deeper coverage enables more comprehensive reconstruction of genome-wide crossover landscapes (Figs **4****, 5**; Table **S1**). However, COmapper does not retain individual-level information, such as the number and position of crossovers per sample, which is essential for quantitative trait locus mapping. Additionally, it cannot be used to assess crossover interference, which requires measuring the physical distances between crossovers along the same chromosome, as can be done with GBS in individually genotyped recombinants (Table **S3**).

In contrast to GBS, COmapper requires a simple gDNA extraction method and pooled gDNA from F_1_ hybrid pollen or F_2_ seedlings for nanopore sequencing, instead of gDNA extraction from individual plants and barcoding. Therefore, we propose that COmapper can be widely used to map genome-wide crossovers at high resolution in the following types of studies. First, COmapper should be applicable for performing a rapid and high-throughput screen to detect changes in the number and distribution of meiotic crossovers in crosses between different accessions in *Arabidopsis* and various species. Therefore, COmapper should allow testing for the effects of genetic, epigenetic and environmental changes on crossover patterning. Second, COmapper should be able to detect crossovers in diverse diploid and polyploid organisms. However, its successful application in complex genomes depends on the SNP density in crossover regions. Therefore, assessing the density and distribution of SNPs across parental genomes is crucial before applying COmapper. For example, we analyzed SNP distribution between two tomato cultivars (*Solanum lycopersicum* and *S. pimpinellifolium*), demonstrating the scalability of COmapper (Fig. **S19**). Ongoing improvements in nanopore sequencing technology such as longer read lengths and lower sequencing error rates will further enhance its utility for large and complex genomes. For polyploids, COmapper can be applied to hybrid autotetraploid *Arabidopsis* plants derived from natural accessions (e.g., Wa-1 × Bla-5) or colchicine-doubled tetraploid (e.g., Col (4x) × L*er* (4x)). Likewise, COmapper could facilitate crossover analysis in allopolyploid plants such as wheat and rapeseed. Third, COmapper should be useful to map genome-wide crossovers in pollen or progeny of outcrossing species, along with calling parental SNPs by nanopore sequencing. Finally, COmapper should allow the measurement of crossover frequency and distribution in meiotic recombination hotspots. Instead of PCR amplification of single ∼5–10-kb molecules from a recombination hotspot (Choi *et al*., 2013), genotyping of pollen or sperm can be performed in pooled samples and combined with nanopore long-read sequencing. Therefore, given the simplicity of sample preparation and the high accuracy in mapping crossovers by long-read nanopore sequencing, we believe that COmapper will be extensively used to understand the mechanisms of meiotic crossover patterning in diverse species beyond plants.

## Acknowledgements

We thank Raphael Mercier (MPI for Plant Breeding Research) for providing *recq4ab* seeds. This work was funded by a Samsung Science and Technology Foundation grant (SSTF-BA2202-09) and the National Research Foundation of Korea (RS-2024-00335818, RS-2024-00407469).

## Author contributions

D.B., N.S. and K.C. designed the project. D.B., N.S., and J.K. developed COmapper pipeline. D.B., N.S., H.K., J.K., J.P., S.P., H.K., J.K., S.L, Y.H., P.A.Z., I.R.H, and K.C. performed experiments and analysis. D.B., N.S. and K.C. wrote the original draft of manuscript. All authors revised the manuscript.

## Data availability

The nanopore sequencing data generated in this study have been deposited in the ArrayExpress database at EMBL-EBL (http://www.ebi.ac.uk/arrayexpress). Detailed information on the nanopore long-read sequencing and other data for GBS and genomic features used in this study is included in **Supplementary Table S2**. Source data are provided with this paper. All custom code used in this study, including the COmapper pipeline, is available at GitHub (https://github.com/KyuhaChoi-Lab/COmapper).

