## Supplementary material for "COmapper: High-resolution mapping of meiotic crossovers by long-read sequencing in *Arabidopsis*": SI.pdf

The following Supporting Information is available for this article:

**Fig. S1** Distribution of read length from long-read nanopore sequencing.

**Fig. S2** Nanopore sequencing read coverage.

**Fig. S3** Comparison of genome structure and single-nucleotide polymorphisms between the Col and L*er* accessions.

**Fig. S4** Validation of COmapper for mapping crossovers using Col × L*er* $F_2$ individuals.

**Fig. S5** Estimating false positive rates attributable to nanopore sequencing errors.

**Fig. S6** Representative simple crossover and complex crossover sites identified by COmapper in nanopore sequencing reads.

**Fig. S7** Representative complex crossover sites identified by both COmapper and GBS allele frequency analysis.

**Fig. S8** Sources of false positives in COmapper results from pooled 4 or 46 Col × L*er* $F_2$ individuals.

**Fig. S9** Analysis of crossovers not detected by COmapper from 46 Col × L*er* $F_2$ individuals.

**Fig. S10** Impact of sequencing coverage on COmapper performance.

**Fig. S11** Validation of COmapper using recq4ab Col × *recq4ab* L*er* $F_2$ recombinant plants.

**Fig. S12** Validation of Compatibility between TAIR10 and Col-CEN Genomes.

**Fig. S13**Comparison of genome-wide crossover landscapes from WT Col × L*er* $F_1$ pollen COmapper replicates.

**Fig. S14** Comparison of SNP distances at crossover sites mapped by GBS and COmapper.

**Fig. S15** Genomic region classification based on 8-SNP length and corresponding crossover distributions.

**Fig. S16** Crossovers occur more frequently at gene promoters and less frequently at gene bodies than a random distribution.

**Fig. S17** Resolution of crossovers mapped by GBS and COmapper.

**Fig. S18** Class I crossovers are associated with higher SNP density.

**Fig. S19** Analysis of SNP density between Solanum *lycopersicum* and Solanum *pimpinellifolium.*

**Table S1** Summary of long-read nanopore sequencing libraries used in this study.

**Table S2** Detailed information for the sequencing libraries used in this study.

**Table S3** Comparison of genome-wide crossover mapping techniques.

**Dataset S1** Allele frequency maps of 46 and 4 F2 individuals.

**Dataset S2** Information of crossover sites in 46 and 4 F2 individuals.

**Dataset S3** Screenshots of false positives.

**Dataset S4** Fisher's exact test for Fig. S9c.

**References**

**Fig. S1 Distribution of read length from long-read nanopore sequencing.**

Distribution of read length from nanopore sequencing of the samples used in this study. Vertical blue lines indicate the N50 read length.

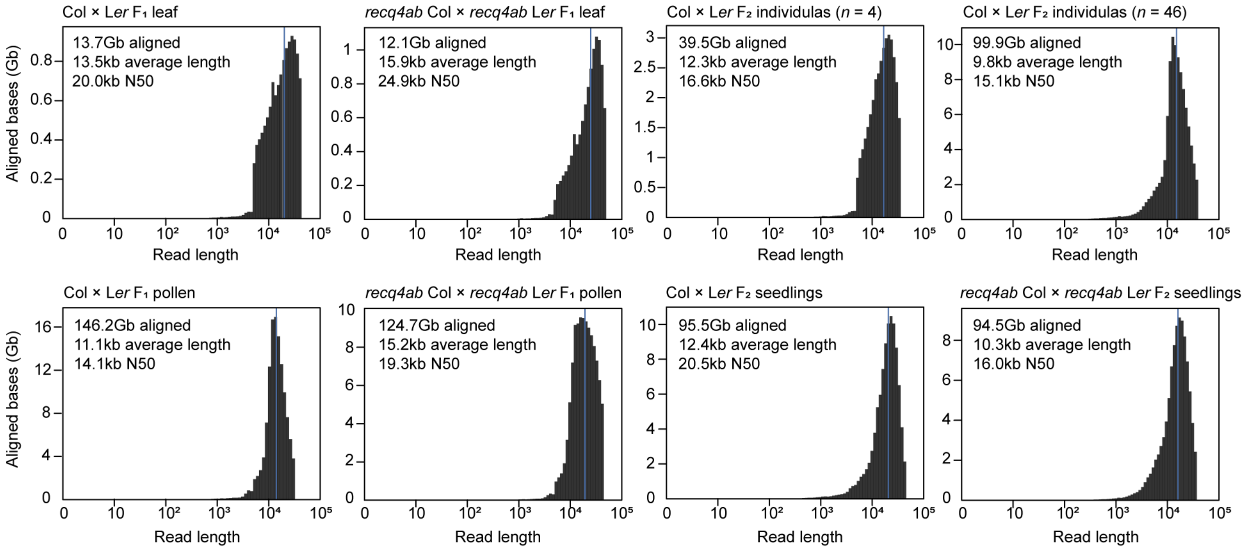

**Fig. S2 Nanopore sequencing read coverage.** Coverage frequencies of long-read nanopore sequencing depth in the number of reads along the five *Arabidopsis* chromosomes on a continuous *x*-axis in the samples used in this study. The vertical dashed lines indicate centromere assembly gap positions, and the vertical lines indicate the ends of the chromosomes. The pericentromeric regions are highlighted in blue.

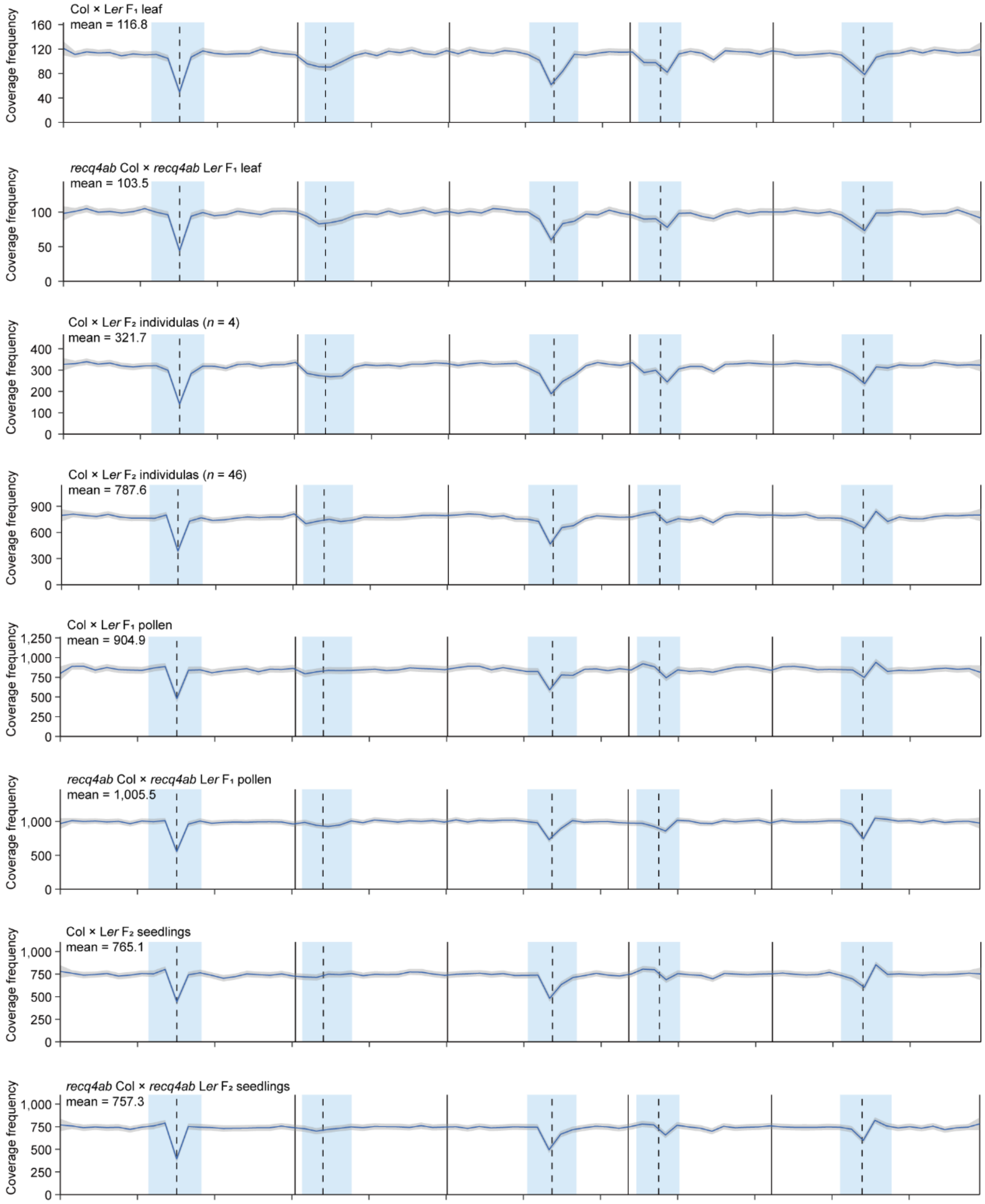

**Fig. S3 Comparison of genome structure and single-nucleotide polymorphisms between the Col and L*er* accessions.** SNP markers (black, SNPs/100 kb), syntenic (gray), translocation (green), duplication (sky blue), inversion (orange) and centromere gaps (black dot) are shown along the five chromosomes of *Arabidopsis* Col (red) and L*er* (blue) accessions.

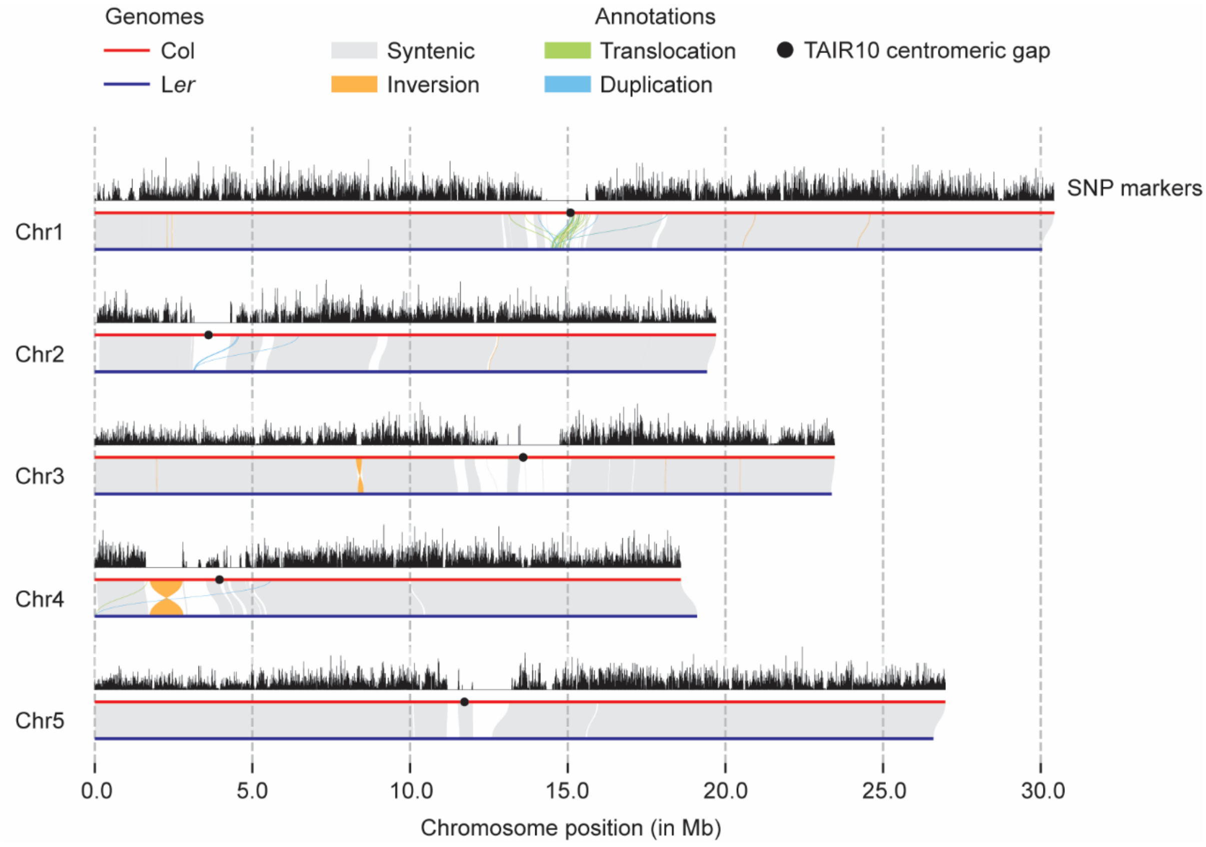

**Fig. S4 Validation of COmapper for mapping crossovers using Col × L*er* $F_2$ individuals.** (**a**) Diagram of the strategy used to map crossovers for each individual by individual barcoding, performing GBS, and running TIGER of genomic DNAs from 4 or 46 Col × L*er* $F_2$ or 39 *recq4ab* Col × *recq4ab* L*er* $F_2$ recombinant individuals. The same genomic DNA purified from each $F_2$ individuals were pooled to generate two separate libraries for nanopore sequencing. (**b**) Summary table of the amount of DNA and genome coverage from long-read nanopore sequencing, and the number of crossovers mapped by the GBS and COmapper approaches.

**(a)**

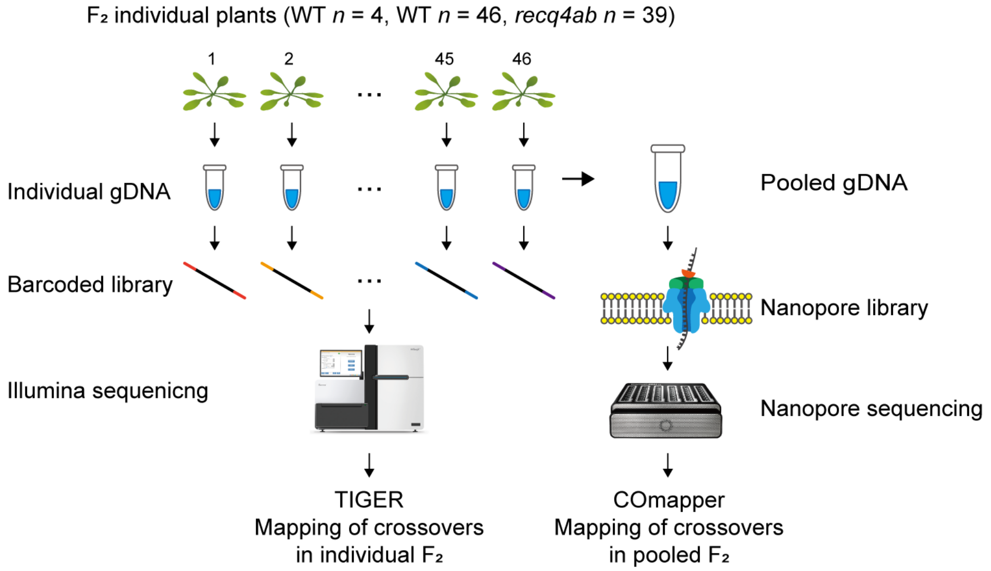


**(b)**

| $F_2$ plants | Reference crossover sites | Nanopore sequenceing coverage (Gb) | Crossover reads mapped by COmapper | Reference sites mapped by COmapper |
|---|---|---|---|---|
| WT (*n* = 4) | 33 | 40.1 | 630 | 33 |
| WT (*n* = 46) | 342 | 100.7 | 1659 | 299 |
| *recq4ab* (*n* = 39) | 1055 | 46.8 | 1448 | 532 |

**Fig. S5 Estimating false positive rates attributable to nanopore sequencing errors.** (**a**) Density plot showing the distribution of average base quality scores and sequence identity of nanopore reads, used to estimate average sequencing error rate. (**b**) Schematic illustration of how consecutive sequencing errors can result in false positive crossovers. The probability of each error was calculated based on a Q14 error rate, assuming that each of the three alternative bases has an equal probability of being erroneously called. The cumulative probability of such errors in the 4-SCO model was estimated to be 0.0357%. (**c**) Table showing the expected and observed numbers of false positive crossovers per million reads across different COmapper models. (**d**) Examples of a false positive crossover identified in the 4-SCO model. Col alleles are represented as 'C', L*er* alleles as 'L', and undetermined alleles as 'N'; 'N' positions were excluded from the COmapper analysis.

(a)

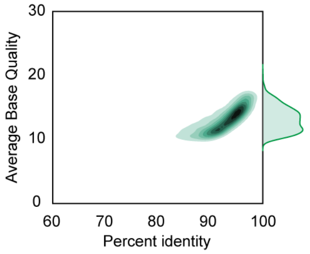


(b)

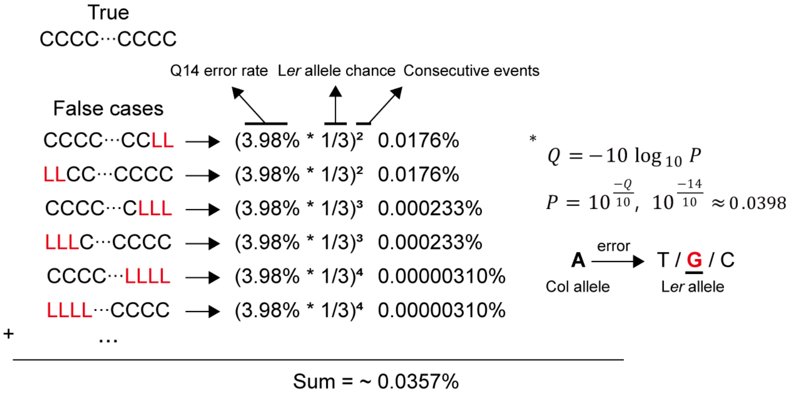


(c)

| | 4-SCO | 6-SCO | 8-SCO | 8-CO |
|---|---|---|---|---|
| Expected | 357 | 5 | 0.06 | 0.12 |
| Observed | 496 | 59 | 3.7 | 3.9 |

(False positives per million reads)

(d)

| | Position | SNP genotypes |
|---|---|---|
| read1 | Chr4::9,264,958-9,286,306 | LNNLLLNLLLNLLLLLLLLNLLLCC |
| read2 | Chr4::16,468,045-16,486,419 | NCCNLLNLLNLLLLLLLNN |
| read3 | Chr5::18,014,967-18,029,258 | CCCNCCCCNCCCCCCLL |
| read4 | Chr1::17,232,360-17,240,240 | LLNCCCCCCCCCC |

**Fig. S6 Representative simple crossover and complex crossover sites identified by COmapper in nanopore sequencing reads.** (**a**) Plots showing the interval of three independent crossovers with a simple exchange between consecutive specific SNPs mapped by COmapper in five nanopore sequencing reads. Red dots indicate Col-specific SNPs, blue dots indicate L*er*-specific SNPs and gray dots indicate missing SNP information due to sequencing errors. Pink areas indicate Col haplotype regions and sky-blue areas indicate L*er* haplotype regions. (**b**) As in (**a**), but showing complex crossover sites.

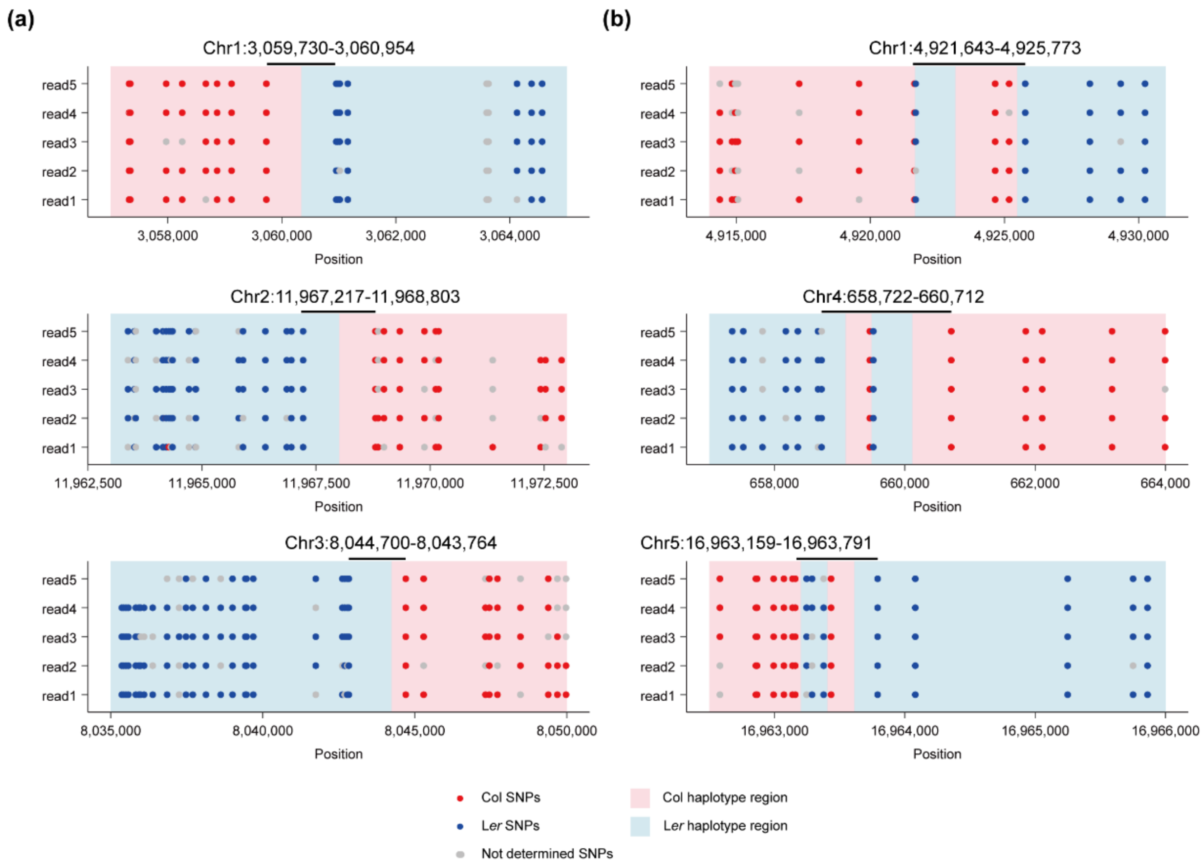

**Fig. S7 Representative complex crossover sites identified by both COmapper and GBS allele frequency analysis.** Plots showing three examples of complex crossover regions. The upper panel displays SNP mapping results from COmapper, while the lower panel shows allele frequency data from GBS. In the upper panel, red dots indicate Col-specific SNPs, blue dots indicate L*er*-specific SNPs, and gray dots indicate missing SNP information due to sequencing errors. In the lower panel, black dots show the L*er* allele frequency based on GBS data, pink areas indicate Col/Col diplotype regions, sky-blue areas indicate Col/L*er* diplotype regions, and yellow triangles point to GBS SNP sites with sequencing depth below 2.

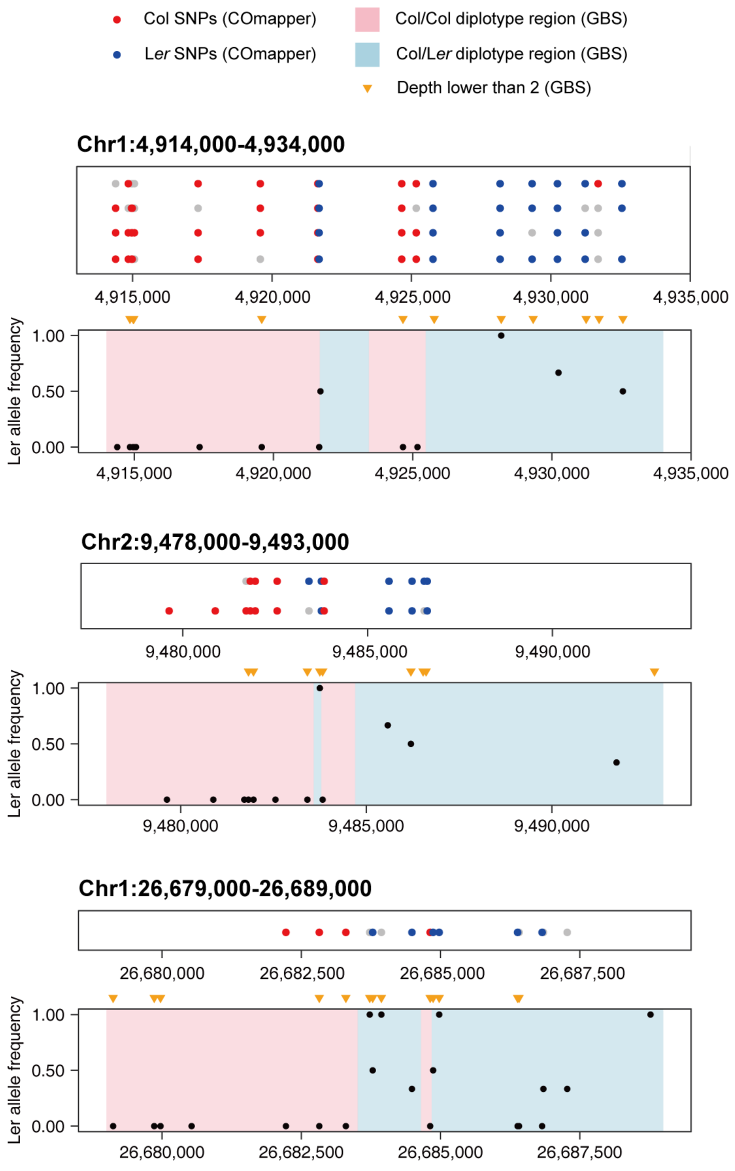

**Fig. S8 Sources of false positives in COmapper results from pooled 4 or 46 Col × L*er* $F_2$ individuals.** The bar plot illustrates the number of different sources of false positive crossovers identified in WT Col × L*er* $F_2$ individuals. (*n* = 4 and *n* = 46). The categories of sources are color-coded as follows: chimeric reads with error clusters (dark blue), substitution errors (orange), mitotic recombination artifacts (green), and chimeric reads with large indels (purple). The total number of false positives observed is indicated above the respective bar.

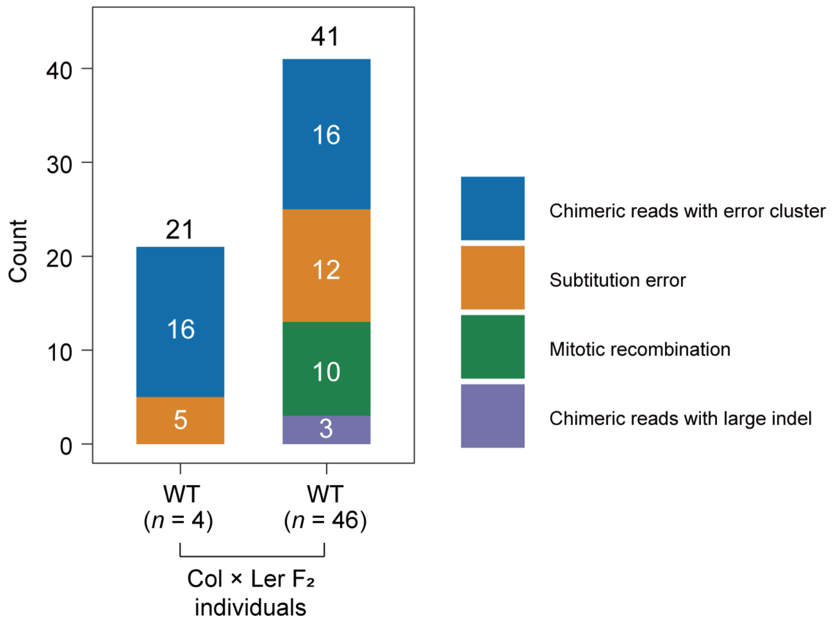

**Fig. S9 Analysis of crossovers not detected by COmapper from 46 Col × L*er* $F_2$ individuals.** (**a**) Number of crossovers detected and undetected by COmapper in the 46-individual pool, categorized by the length of 8 SNPs flanking the crossover sites. (**b**) Proportion of detected and undetected crossovers across the same 8-SNP length categories as in (a). (**c**) Genomic distribution of detected (blue) and undetected (red) crossovers along the five chromosomes (TAIR10 reference genome). Vertical dotted lines mark the centromeric gaps; grey regions indicate pericentromeric regions.

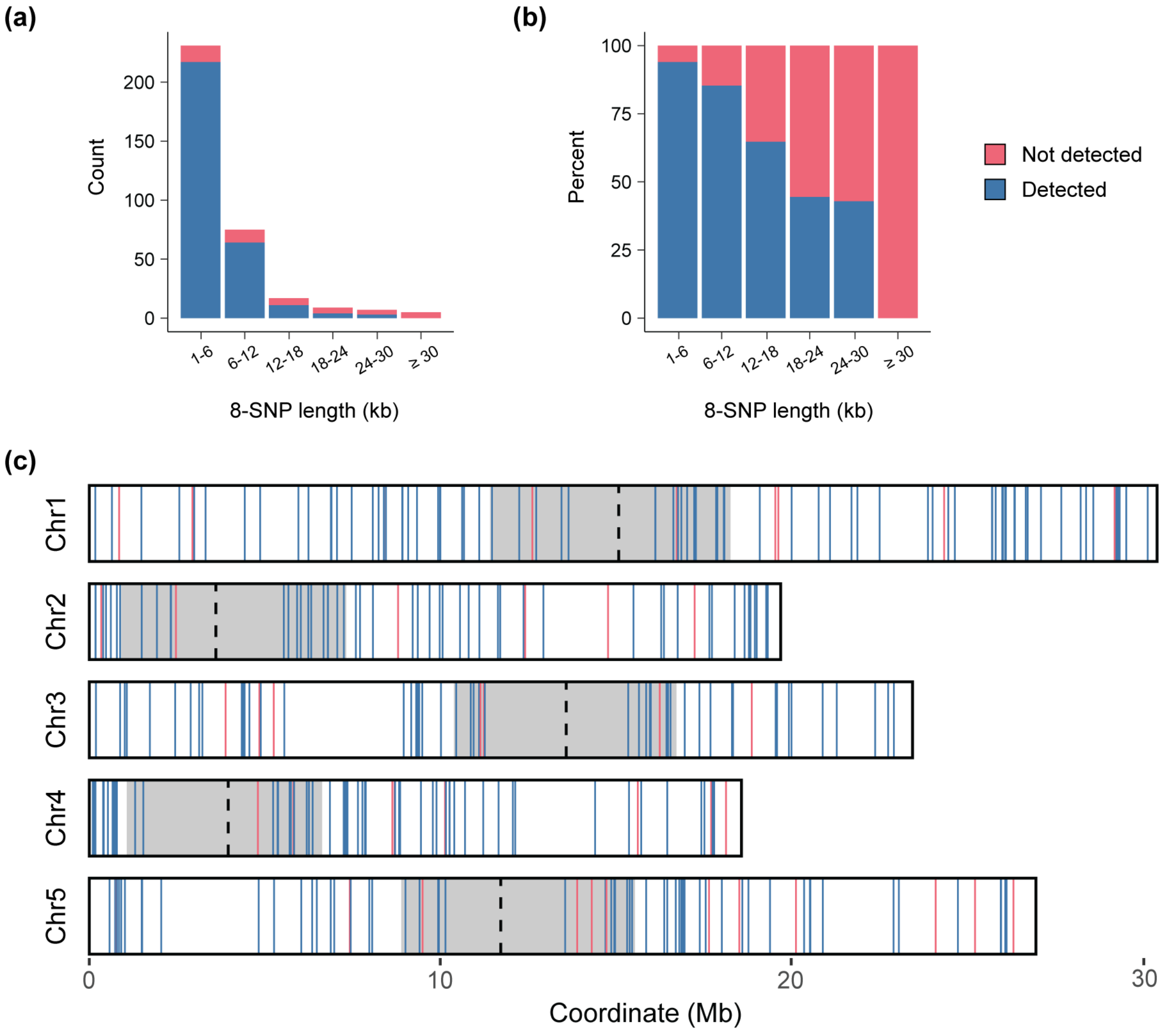

**Fig. S10 Impact of sequencing coverage on COmapper performance.** (**a-c**) True positive (**a**), false positive (**b**), and precision (TP/[TP + FP]) (**c**) as a function of sequencing coverage, based on subsampling of data from for 4 $F_2$ individuals. (**d-f**) Same as (**a-c**), but based on subsampling of data from 46 $F_2$ individuals. (**g**) Number of undetected crossovers (false negatives), plotted as a function of coverage per $F_2$. (**h**) The proportion of detected crossovers relative to total crossovers in the 4 $F_2$ dataset, plotted as a function of sequencing coverage per $F_2$. Vertical lines represent coverage per $F_2$ for 4 (blue), 46 (red) $F_2$ individuals, and 1,000 (yellow) $F_2$ seedlings. (**i**) Proportion of non-overlapping crossover sites per total detected crossovers, as a function of sequencing coverage per $F_2$. Vertical lines indicate coverage per $F_2$ for 4 (blue), 46 (red) $F_2$ individuals, and 1,000 (yellow) $F_2$ seedlings.

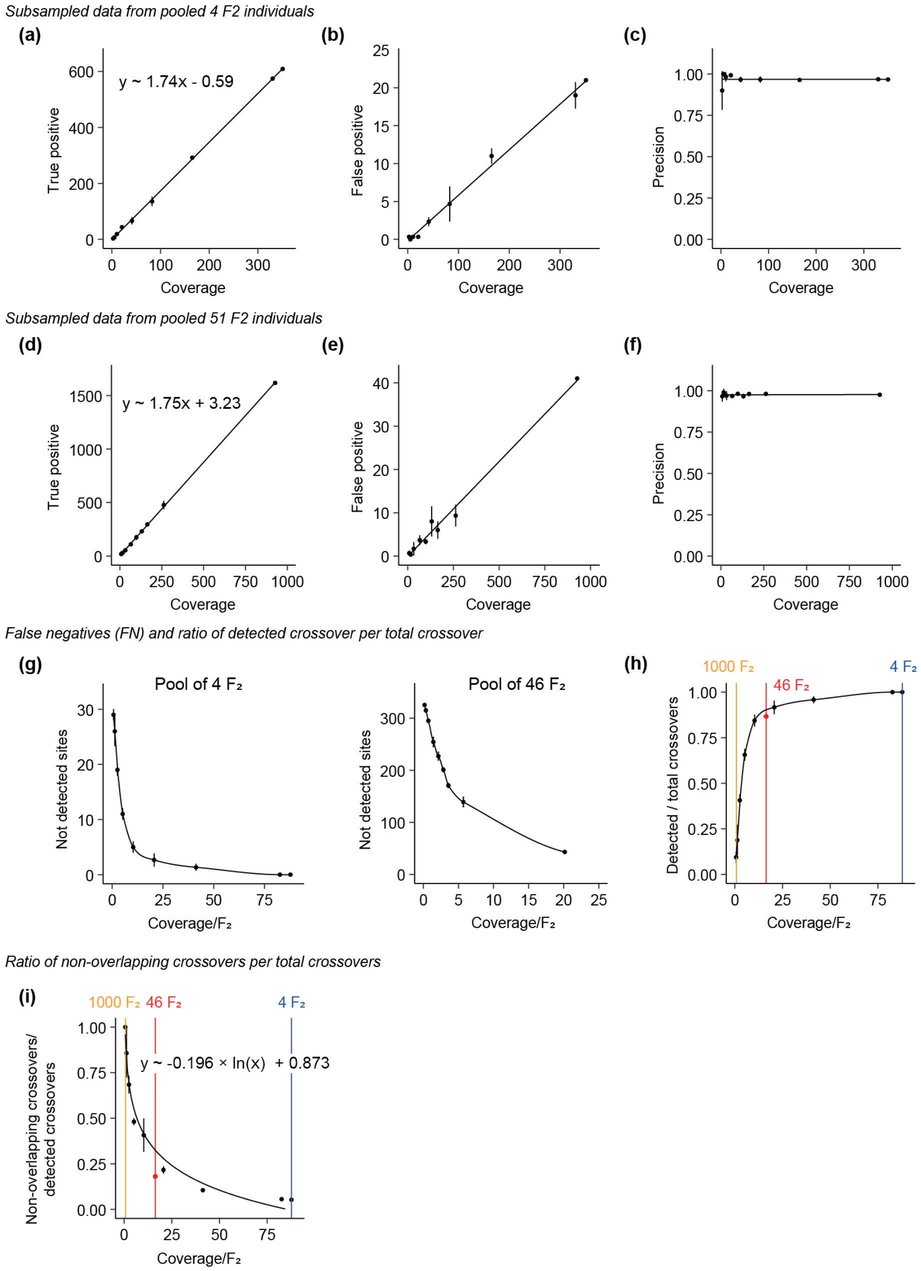

**Fig. S11 Validation of COmapper using recq4ab Col × *recq4ab* L*er* $F_2$ recombinant plants.** (**a**) Number of crossovers detected per million reads in different COmapper models. True crossovers (blue) were detected by COmapper models among the crossovers mapped by GBS in 39 *recq4ab* Col × *recq4ab* L*er* $F_2$ plants, and false crossovers (red) indicate the crossovers that differ from the crossovers defined by GBS. (**b**) Precision of crossover detection in the different COmapper models, include Col × L*er* $F_2$ plants shown in Fig. **3e**. (**c**) Precision of 8-CO COmapper model in WT and *recq4ab* $F_2$ individuals, respectively done by random subsampling. (**d**) Crossover per haploid in Col-0 × L*er* $F_2$ individuals and *recq4ab* Col × *recq4ab* L*er* $F_2$ individuals. (**e**) Relative proportion of crossover types of *recq4ab* Col × *recq4ab* L*er* $F_2$ individuals with Col × L*er* $F_2$ individuals, $F_1$ pollen, and $F_2$ seedlings data shown in Fig. **3f**. (**f**) Table of crossover numbers, coverage, and precision of Col × L*er* $F_2$ individuals (*n* = 4 and *n* = 46) and *recq4ab* Col × *recq4ab* L*er* $F_2$ individuals (*n* = 39). (**g**) Categorization of false positive sources in *recq4ab* Col × *recq4ab* L*er* $F_2$ individuals with Col × L*er* $F_2$ false positive sources data shown in Fig. **S8**.

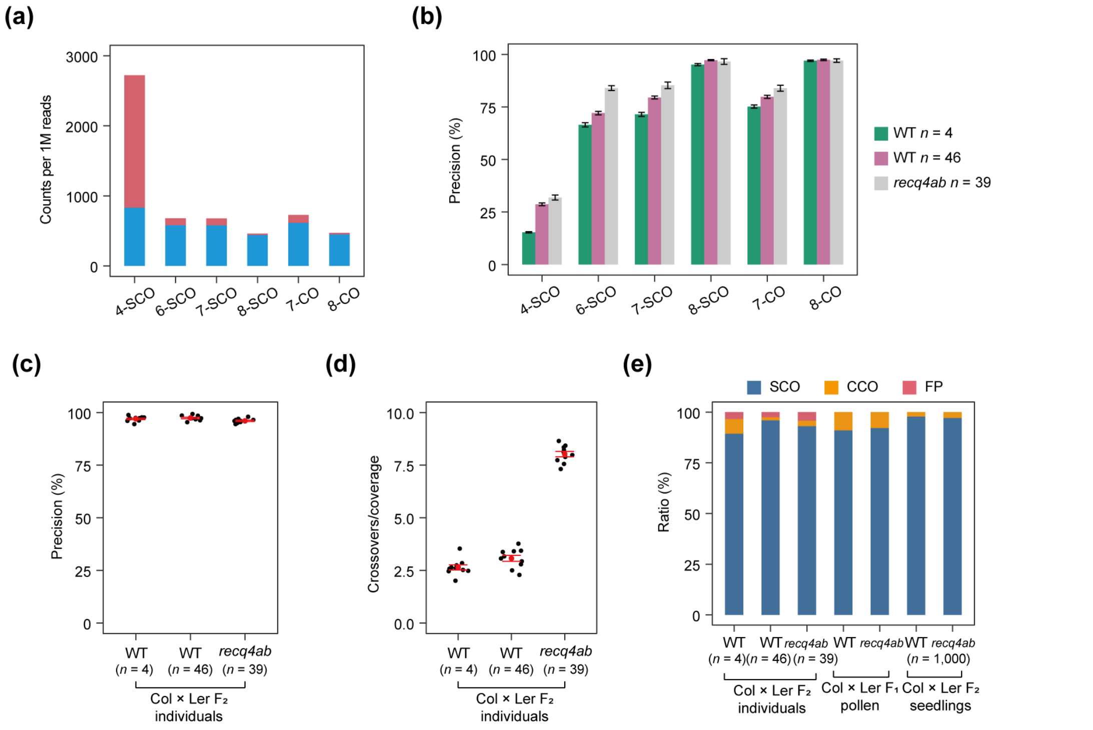


(f)

| | n | Reference crossovers (GBS) | Coverage (COmapper) | True positive | False positive | Not detected | Precision (%) |
|---|---|---|---|---|---|---|---|
| WT $F_2$ | 4 | 33 | 337 | 609 | 21 | 0 | 96.7 |
| WT $F_2$ | 46 | 342 | 845 | 1618 | 41 | 45 | 97.5 |
| recq4 $F_2$ | 39 | 1055 | 410 | 1387 | 64 | 523 | 95.6 |

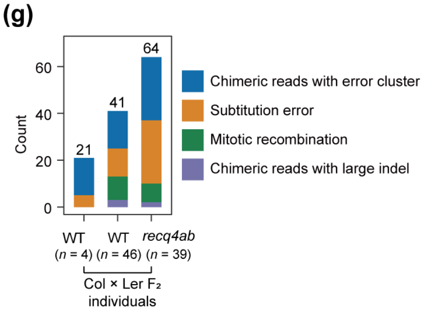

**Fig. S12 Validation of Compatibility between TAIR10 and Col-CEN Genomes.** (**a**) Procedure for TAIR10 SNP converting to Col-CEN using Liftoff pipeline. (**b**) A Venn diagram illustrating the overlap of crossover reads mapped by TAIR10 and Col-CEN reference genome within a set of 46 Col-0 × L*er* $F_2$ individuals. The overlapping region (purple) represents 1,658 SNPs that were consistently identified and mapped to both reference genomes. The non-overlapping regions (light red and light blue) indicate SNPs uniquely mapped to either TAIR10 or Col-CEN.

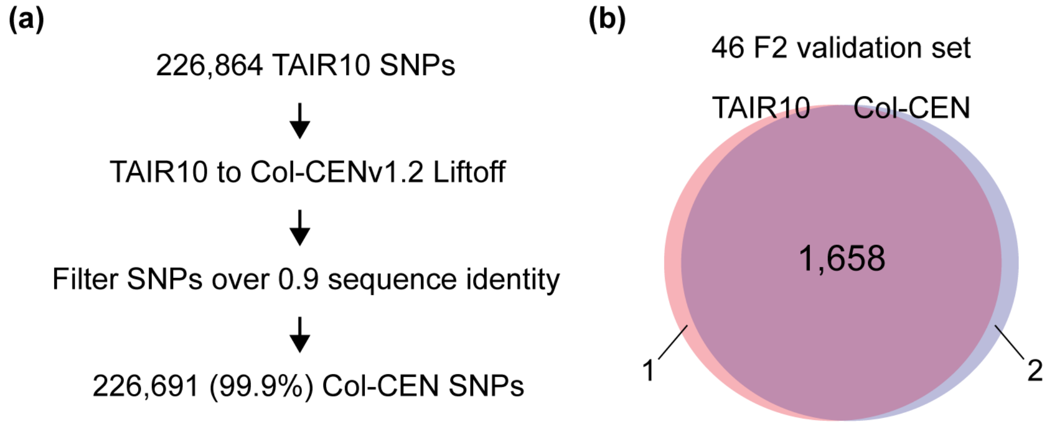

**Fig. S13 Comparison of genome-wide crossover landscapes from WT Col × L*er* $F_1$ pollen COmapper replicates.** (**a**) Genome-wide crossover landscapes obtained from two independent WT Col × L*er* $F_1$ pollen COmapper experiments. Relative crossover (CO) rates were calculated in non-overlapping 100-kb bins by dividing the number of crossovers per bin by the total number of crossovers, followed by smoothing using moving average. (**b**) As in (**a**), but comparing crossover landscape from GBS of WT Col × L*er* $F_1$ male meiosis with WT Col × L*er* $F_1$ pollen COmapper replicate 1. (**c**) As in (**a**), but comparing crossover landscape from GBS of WT Col × L*er* $F_1$ male meiosis with WT Col × L*er* $F_1$ pollen COmapper set 2. Fisher's exact test was applied to each 100-kb bin to assess differences in relative crossover rates, with P-values corrected using Benjamini-Hochberg method. Bins with false discovery rate (FDR) < 0.05 are highlighted in yellow.

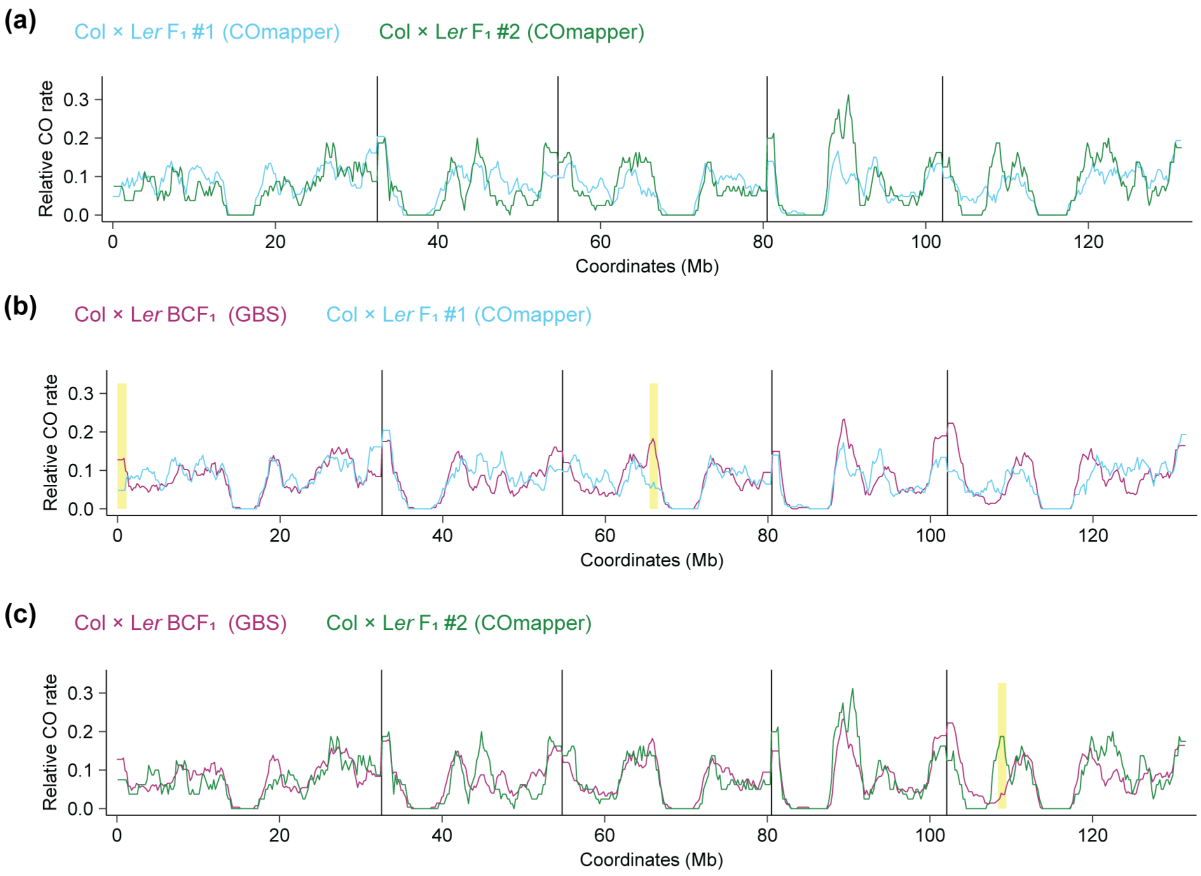

**Fig. S14 Comparison of SNP distances at crossover sites mapped by GBS and COmapper.** (**a**) Relative crossovers mapped by GBS (blue, 1,824 crossovers) and COmapper (red, 1,679 crossovers) along the chromosome telomere (TEL)-to-centromere (CEN) axis. (**b**) Summary of the physical coordinates of sub-telomeric, mid-chromosomal, pericentromeric, and centromeric regions for each chromosome. (**c**) SNP density in sub-telomeric, mid-chromosomal and pericentromeric regions along chromosomes. Vertical dashed lines indicate the median number of SNPs per 10-kb window. Asterisks indicate significant differences between regions (**P* < 0.05, ***P* < 0.01, ****P* < 0.001; Wilcoxon test). (**d**) Histogram showing the length of eight-SNP strings at crossovers mapped by GBS (blue, Col × L*er* $BCF_1$) and COmapper (red, Col × L*er* $F_1$ pollen) for male meiosis in sub-telomeric regions. Vertical dashed lines indicate the median length (in bp) of eight-SNP strings. Asterisks indicate significant differences between genotypes (**P* < 0.05, ***P* < 0.01, ****P* < 0.001; Wilcoxon test). (**e**) As in (**d**), but showing the mid-chromosomal regions. (**f**) As in (**d**), but showing the pericentromeric regions.

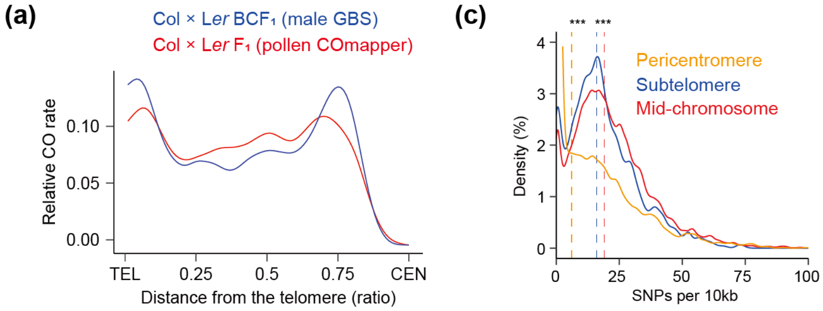


(b)

| Chr. | Subtelomere (northern arm) | Mid-chromosome (northern arm) | Pericentromere | Mid-chromosome (southern arm) | Subtelomere (southern arm) |
|---|---|---|---|---|---|
| 1 | 1~1,508,604 | 1,508,605~ 11,420,000 | 11,420,001~ 18,270,000 | 18,270,001~ 28,893,508 | 28,893,509~ 30,427,671 |
| 2 | 1~910,000 | | 910,001~ 7,320,000 | 7,320,001~ 18,089,253 | 18,089,254~ 19,698,289 |
| 3 | 1~1,358,778 | 1,358,779~ 10,390,000 | 10,390,001~ 16,730,000 | 16,730,001~ 22,472,625 | 22,472,626~ 23,459,830 |
| 4 | 1~1,070,000 | | 1,070,001~ 6,630,000 | 6,630,001~ 17,122,152 | 17,122,153~ 18,585,056 |
| 5 | 1~1,172,502 | 1,172,503~ 8,890,000 | 8,890,001~ 15,550,000 | 15,550,001~ 25,450,454 | 25,450,455~ 26,975,502 |

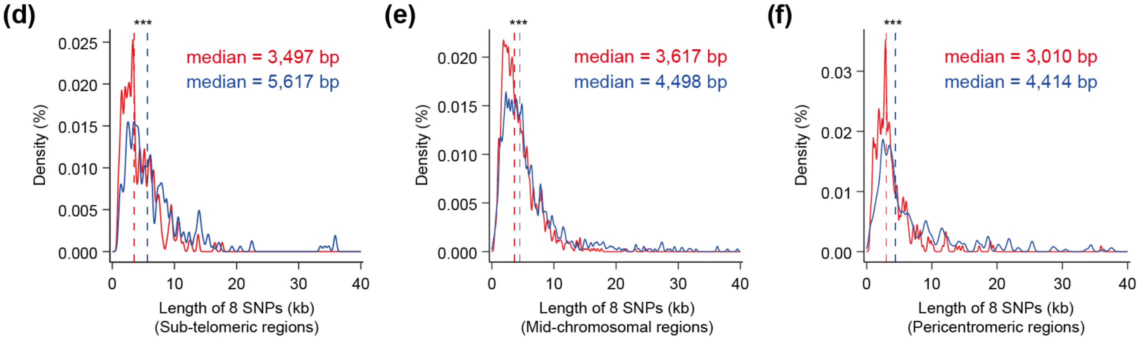

**Fig. S15 Genomic region classification based on 8-SNP length and corresponding crossover distributions.** (**a**) Genomic regions were classified into four categories based on the length of 8 SNPs flanking each SNP interval. (**b**) The proportion of the genome corresponding to each 8-SNP length category is shown. The proportion of regions with 8-SNP lengths ≥ 15 kb that overlap with *CEN178* repeats or structural variants (SVs) is also indicated. (**c**) Proportions of crossovers occurring within each genomic category. Crossovers detected by GBS and COmapper (from Col × L*er* $F_2$ and $F_1$ pollen) are shown separately.

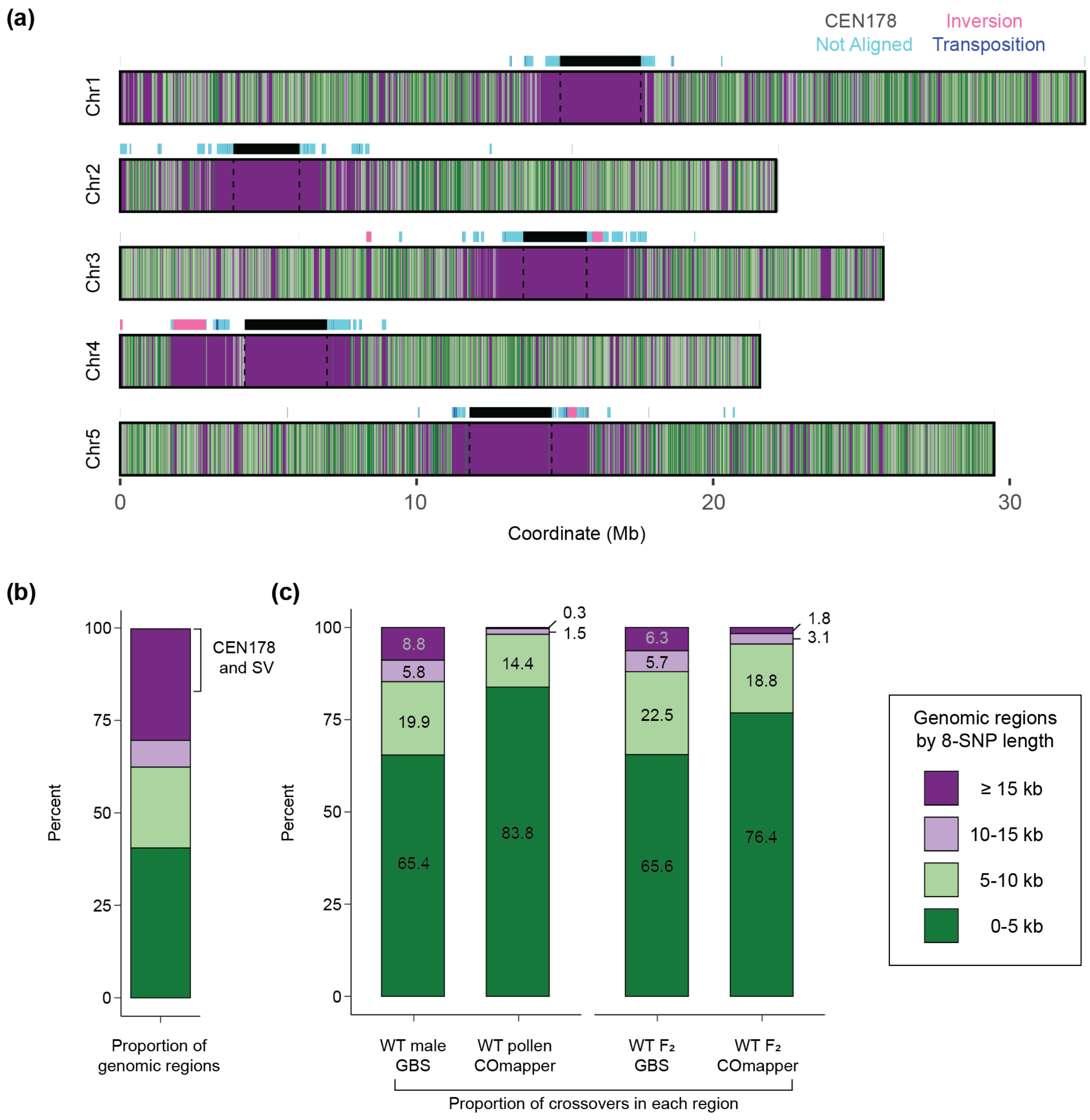

**Fig. S16 Crossovers occur more frequently at gene promoters and less frequently at gene bodies than a random distribution.** (**a**) Diagram showing the number of 0.5-kb bins around gene transcription start sites (TSS) and end sites (TES) in a 4-kb window for the analysis of crossovers. (**b**) Number of crossovers in each bin for crossovers mapped by GBS (left, Col × L*er* BCF$_1$ plants) or COmapper (right, Col × L*er* F$_1$ pollen) and number of crossovers from a random distribution. Asterisks indicate significant differences compared to the random distribution (**P* < 0.05, ***P* < 0.01, ****P* < 0.001; Wilcoxon test). (**c**) As in (**b**), but showing data for Col × L*er* F$_2$ plants (GBS) and Col × L*er* F$_2$ seedlings (COmapper). (**d**) As in (**c**), but showing data for *recq4ab* Col × *recq4ab* L*er* F$_2$ plants (GBS) and seedlings (COmapper).

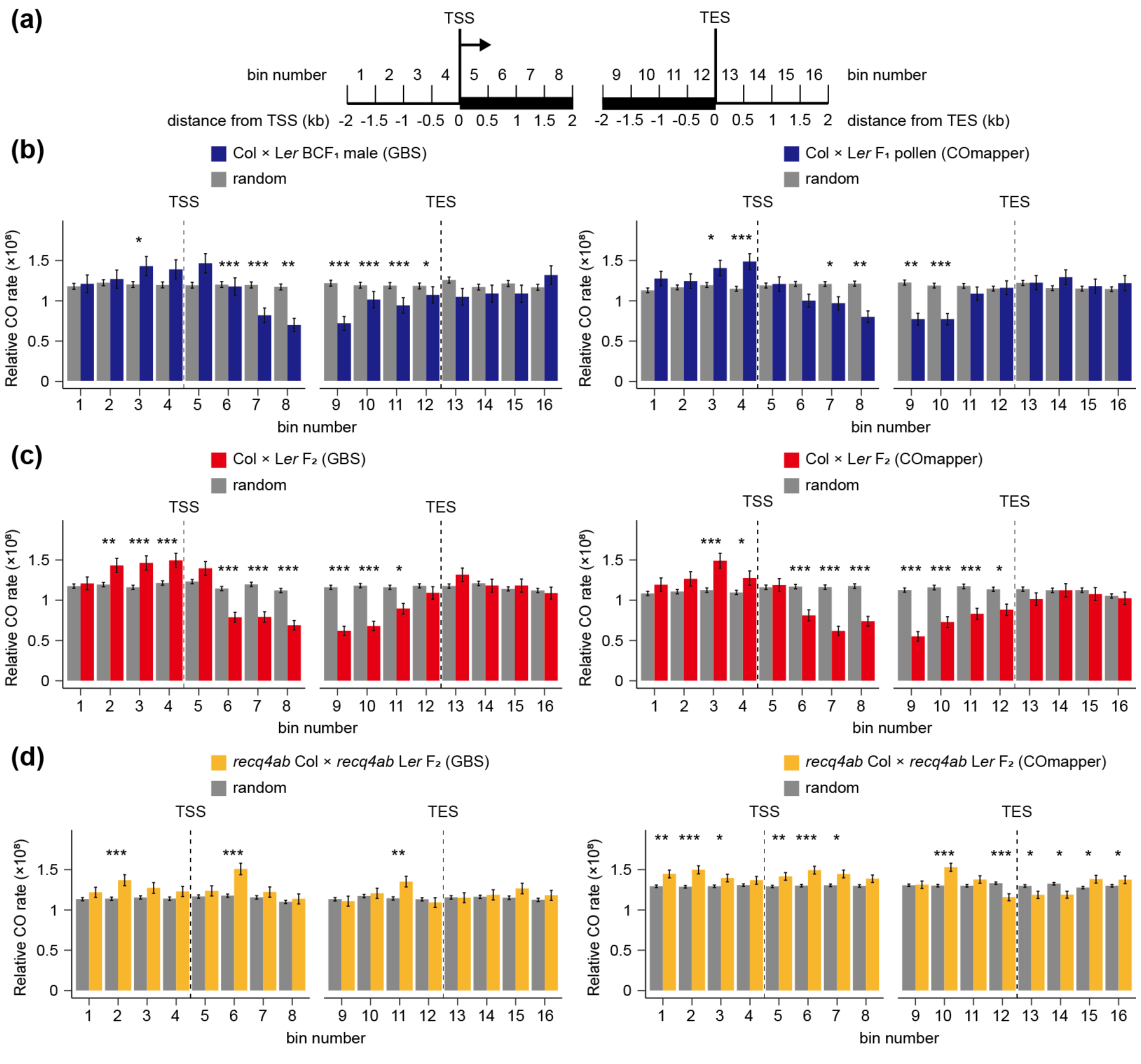

**Fig. S17 Resolution of crossovers mapped by GBS and COmapper.** (**a**) A model showing the resolution of crossover sites detected by COmapper and GBS. The resolution refers to the distance between the genotyped SNPs flanking the crossover site. COmapper directly determines the crossover site and the resolution is defined as the distance between the closest two SNPs surrounding it. In contrast, the resolution of crossover sites detected by GBS decreases with reduced library sequencing coverage around crossover sites. (**b**) Boxplot comparing resolution of crossovers grouped by sequencing coverage of GBS libraries. Crossover resolution from COmapper of Col × L*er* $F_2$ seedling and Col × L*er* $F_1$ pollen is shown. Different lowercase letters indicate significant differences ($P < 0.05$) as determined by ANOVA followed by the Games-Howell test. (**c**) Statistical analysis for resolution of crossovers mapped by GBS and COmapper, with GBS libraries grouped by coverage. SD = standard deviation. SE = standard error. (**d**) Metaplots showing the density of crossovers mapped by GBS. GBS libraries were divided into three coverage groups for plotting.

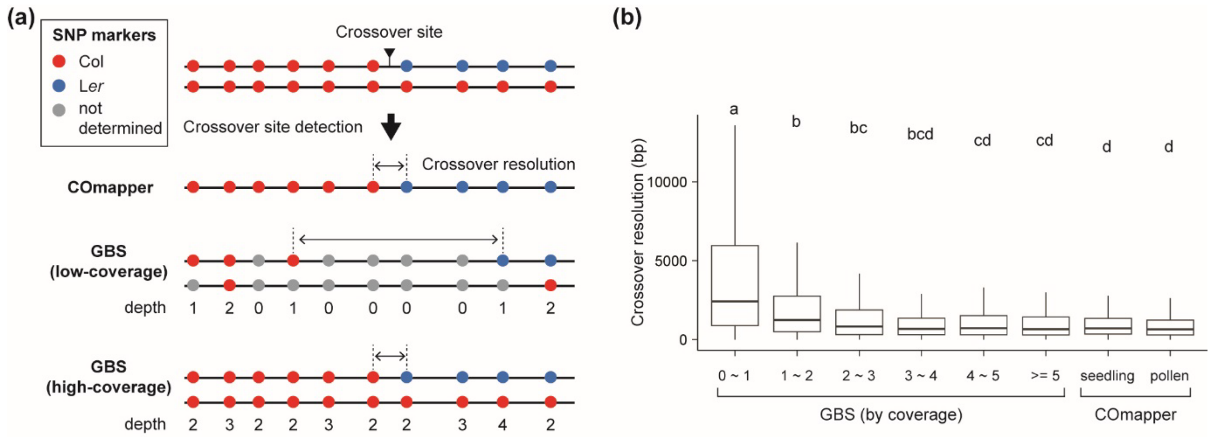


(c)

| Method | Group | Libraries | Crossovers | Resolution (bp) | | | |
|---|---|---|---|---|---|---|---|
| | | | | Median | Mean | SD | SE |
| GBS | 0 - 1x | 777 | 6,313 | 2,422 | 6,106 | 14,329 | 180 |
| | 1 - 2x | 971 | 7,839 | 1,247 | 3,013 | 8,208 | 93 |
| | 2 - 3x | 115 | 948 | 830 | 2,293 | 8,413 | 274 |
| | 3 - 4x | 85 | 696 | 684 | 2,289 | 10,183 | 386 |
| | 4 - 5x | 66 | 557 | 729 | 1,921 | 6,161 | 261 |
| | >= 5x | 81 | 577 | 667 | 1,835 | 5,635 | 235 |
| COmapper | $F_2$ seedling | 1 | 1,433 | 716 | 1,265 | 2,164 | 57 |
| | $F_1$ pollen | 1 | 1,246 | 658 | 1,054 | 1,466 | 42 |

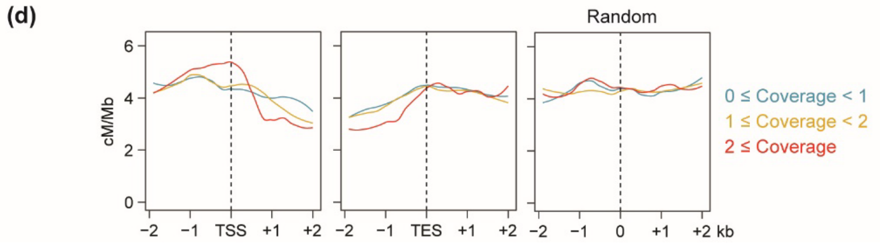

**Fig. S18 Class I crossovers are associated with higher SNP density.** (**a**) Metaplots showing that the crossovers from $F_2$ plants derived from a *J3pro:J3*$^{G155R}$ Col × L*er* cross are associated with higher SNP density in a 2-kb window compared to a random distribution, as shown in WT Col × L*er* hybrid $F_2$ plants (Fig. **6a**). *n* indicates the number of crossover sites mapped by GBS or the number of randomly chosen regions. The left plot shows SNP density around crossovers mapped by GBS compared to random (middle plot). The right plot shows the significance of changes in SNP density according to the physical distance (bp) from the crossover site. Black, green and red dots indicate SNP density per 40-bp bin. Significance between genotype and the random distribution per 40-bp bin was tested using a Wilcoxon test. Red and blue dots above the plot indicate higher and lower SNP density than a random distribution, respectively, while gray dots indicate non-significance. (**b**) As in (**a**), but showing the crossovers mapped by GBS in $F_2$ individual plants from self-pollinated Col × Di-G $F_1$ hybrids.

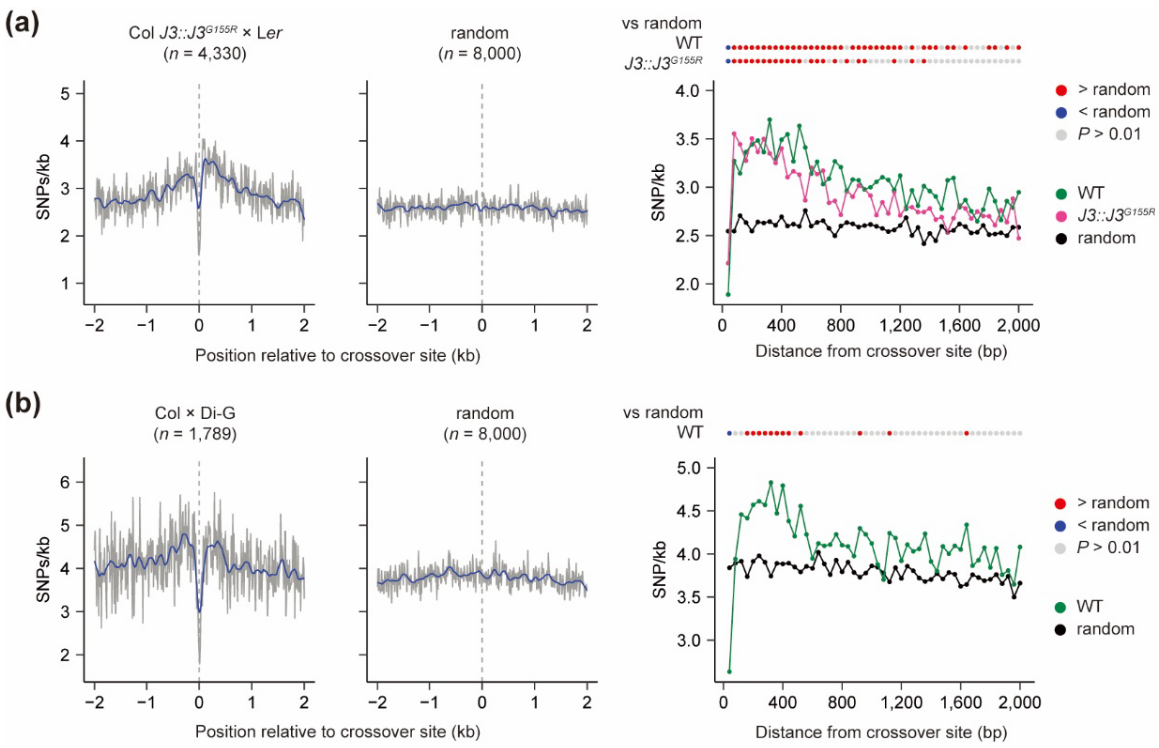

**Fig. S19 Analysis of SNP density between Solanum *lycopersicum* and Solanum *pimpinellifolium.*** (**a**) Genomic regions were classified into four categories based on the length of 8 SNPs flanking each SNP interval. (**b**) The proportion of the genome corresponding to each 8-SNP length category is shown.

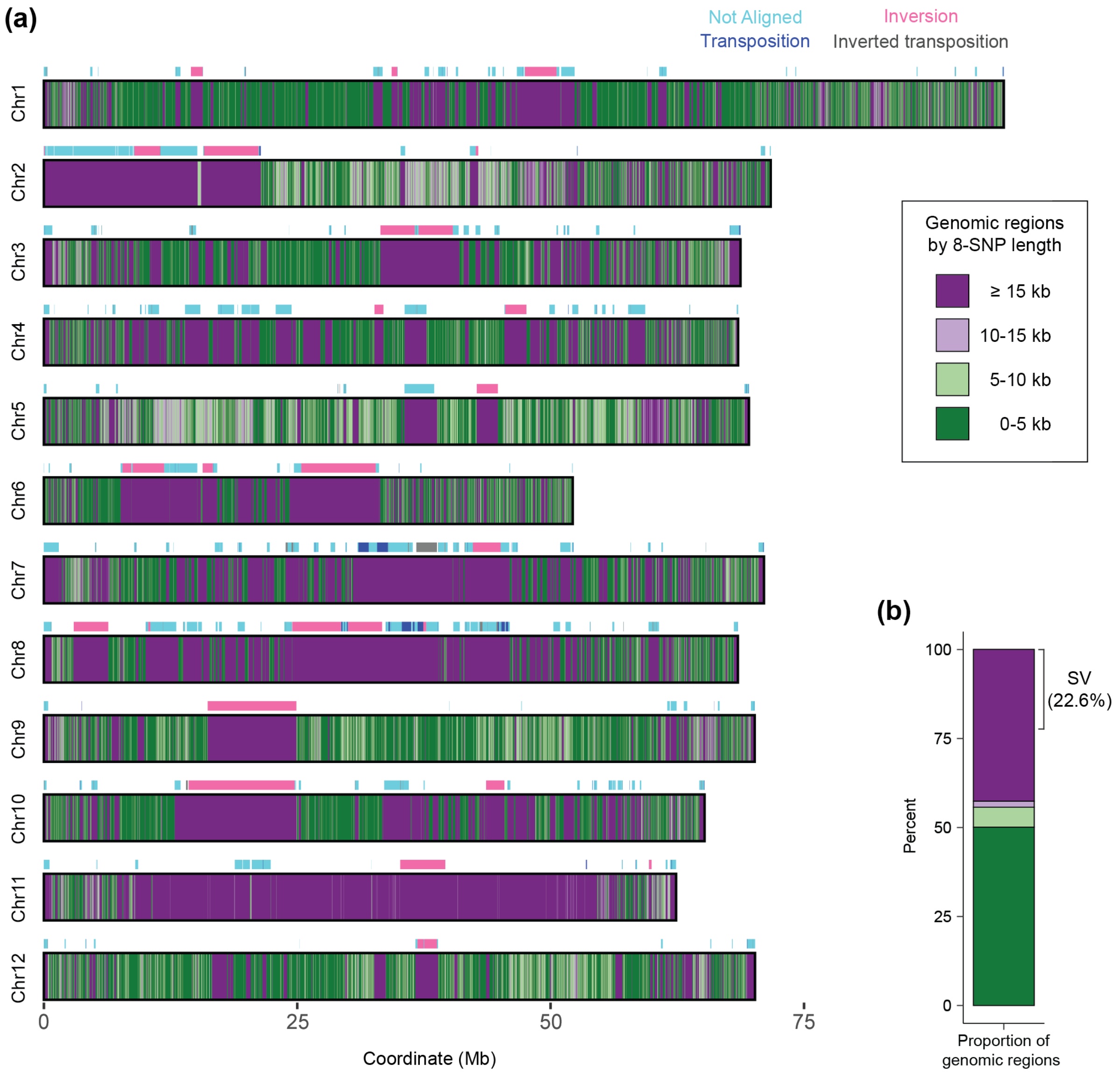

**Table S1. Summary of long-read nanopore sequencing libraries used in this study.** Ten samples with different genotypes and tissues were used to generate long-read nanopore sequencing libraries using high-molecular-weight genomic DNA. *n* indicates the number of pooled Col × L*er* $F_2$ individual plants. The information for the crossover analysis by COmapper is provided for each library.

| Genotype Tissue | Aligned reads | Aligned bases (Gb) | Crossover reads (allows overlap) | Parental reads | Crossovers /haploid |
|---|---|---|---|---|---|
| Col × L*er* $F_1$ leaves | 1,095,831 | 13.99 | 2 | 561,814 | 0.03 |
| *recq4ab* Col × *recq4ab* L*er* $F_1$ leaves | 817,791 | 12.33 | 0 | 436,702 | 0.00 |
| Col × L*er* $F_2$ leaves (*n* = 4) | 3,429,425 | 40.14 | 630 | 1,772,364 | 2.79 |
| Col × L*er* $F_2$ leaves (*n* = 46) | 11,345,385 | 100.74 | 1,659 | 4,895,997 | 2.99 |
| *recq4ab* Col × *recq4ab* L*er* $F_2$ leaves (*n* = 39) | 9,747,722 | 48.83 | 1,448 | 3,546,114 | 7.87 |
| Col × L*er* $F_1$ pollen set1 | 14,041,724 | 149.34 | 1,146 | 9,773,260 | 1.59 |
| Col × L*er* $F_1$ pollen set2 | 4,557,585 | 61.98 | 533 | 4,294,489 | 1.70 |
| *recq4ab* Col × *recq4ab* L*er* $F_1$ pollen | 8,789,632 | 126.66 | 1,963 | 4,988,761 | 3.92 |
| Col × L*er* $F_2$ seedlings (*n* = 1000) | 8,433,348 | 96.14 | 1,679 | 3,553,262 | 3.22 |
| *recq4ab* Col × *recq4ab* L*er* $F_1$ seedlings (*n* = 1000) | 9,980,173 | 95.35 | 6,719 | 4,685,186 | 12.42 |

**Table S2. Detailed information for the sequencing libraries used in this study.** The short-read next-generation and long-read nanopore sequencing libraries used in this study are listed and their information is provided. All data used in this study have been deposited in the ArrayExpress database at EMBL-EBL (http://www.ebi.ac.uk/arrayexpress).

| Library | Dataset accession | Genotype | Read length | Tissue | References |
|---|---|---|---|---|---|
| gDNA | E-MTAB-6257 | Col | 2×100 bp | Floral buds | (Choi et al. 2018) |
| SPO11-1–oligo | E-MTAB-5041 | Col | 2×100 bp | Floral buds | (Choi et al. 2018) |
| MNase-seq | E-MTAB-5042 | Col | 2×100 bp | Floral buds | (Choi et al. 2018) |
| GBS | E-MTAB-8165<br>E-MTAB-10168<br>E-MTAB-10657 | Col × L*er* | 2×150 bp | Leaves | (Rowan *et al.*, 2019; Nageswaran *et al.*, 2021; Kim *et al.*, 2022, 2024) |
| GBS | E-MTAB-12726 | *recq4ab* Col × *recq4ab* L*er* | 2×150 bp | $F_2$ leaves | (Kim *et al.*, 2024) |
| GBS | E-MTAB-13414 | *J3pro:J3$^{G155R}$* Col × L*er* | 2×150 bp | $F_2$ leaves | (Kim *et al.*, 2024) |
| GBS | E-MTAB-14382 | Col × Di-G | 2×150 bp | $F_2$ leaves | This study |
| GBS | E-MTAB-14380 | Col × L*er* | 2×150 bp | 55 $F_2$ leaves | This study |
| GBS | E-MTAB-12695<br>E-MTAB-15084 | L*er* × Col/L*er* | 2×150 bp | $BCF_1$ leaves | This study |
| GBS | E-MTAB-15081 | *recq4ab* L*er* × *recq4ab* Col/L*er* | 2×150 bp | $BCF_1$ leaves | This study |
| Nanopore-seq | E-MTAB-14381 | Col × L*er* | ~13.5 kb | $F_1$ leaves | This study |
| Nanopore-seq | E-MTAB-14381 | *recq4ab* Col × *recq4ab* L*er* | ~15.9 kb | $F_1$ leaves | This study |
| Nanopore-seq | E-MTAB-14367 | Col × L*er* | ~12.3 kb | 4 $F_2$ leaves | This study |
| Nanopore-seq | E-MTAB-14367 | Col × L*er* | ~9.8 kb | 46 $F_2$ leaves | This study |
| Nanopore-seq | E-MTAB-15082 | *recq4ab* Col × *recq4ab* L*er* | ~9.1 kb | 39 $F_2$ leaves | This study |
| Nanopore-seq | E-MTAB-14369 | Col × L*er* | ~14.1 kb | $F_1$ pollen | This study |
| Nanopore-seq | E-MTAB-15083 | Col × L*er* | ~6.8 kb | $F_1$ pollen | This study |
| Nanopore-seq | E-MTAB-14369 | *recq4ab* Col × *recq4ab* L*er* | ~19.3 kb | $F_1$ pollen | This study |
| Nanopore-seq | E-MTAB-14368 | Col × L*er* | ~20.5 kb | $F_2$ seedlings | This study |
| Nanopore-seq | E-MTAB-14368 | *recq4ab* Col × *recq4ab* L*er* | ~16.0 kb | $F_2$ seedling | This study |

**Table S3. Comparison of genome-wide crossover mapping techniques.** Summarizes key features of four genome-wide crossover (CO) mapping techniques: GBS, linked-read sequencing, Hi-C, and COmapper. Each method differs in detection principle, resolution, and data output.

| Feature | GBS | Linked-read-seq | Hi-C | COmapper |
|---|---|---|---|---|
| Detection Principle | - Short read seq<br>- $F_2$ individuals | - Short read seq<br>- Reconstruct long DNA<br>- 10X index<br>- Sample pool | - Shor read seq<br>- Hi-C fragments<br>- Read-pairs<br>- Sample pool | - Long read seq<br>- Sample pool |
| Crossover Detection | - Indirect<br>- Chromosome | - Indirect<br>- Large molecule | - Indirect<br>- Proximity-ligated molecule | - Direct<br>- Long read |
| Resolution | - High resolution with high coverage | - High resolution with high coverage | - Low resolution | - High resolution<br>- Single long read |
| Advantages | - Available on high/low coverage<br>- Individual information<br>- Low SNP density | - High resolution<br>- Pooled sample<br>- Low SNP density | - Chromosome structure<br>- Low SNP density | - High resolution<br>- Pooled sample<br>- Simple library prep |
| Disadvantages | - Individual index<br>- Difficult library prep | - Difficult library prep<br>- Missing individual data | - Low resolution<br>- Difficult library prep<br>- Missing individual data | - Missing individual data<br>- High SNP density |
